# Loss-of-function mutation in human *Oxidation Resistance gene 1* disrupts the spatial-temporal regulation of histone arginine methylation in early brain development

**DOI:** 10.1101/2022.05.24.493324

**Authors:** Xiaolin Lin, Wei Wang, Mingyi Yang, Nadirah Damseh, Mirta Mittelstedt Leal de Sousa, Fadi Jacob, Anna Lång, Elise Kristiansen, Marco Pannone, Miroslava Kissova, Runar Almaas, Anna Kuśnierczyk, Richard Siller, Maher Shahrour, Motee Al-Ashhab, Bassam Abu-Libdeh, Wannan Tang, Geir Slupphaug, Orly Elpeleg, Stig Ove Bøe, Lars Eide, Gareth J Sullivan, Johanne Egge Rinholm, Hongjun Song, Guo-li Ming, Barbara van Loon, Simon Edvardson, Jing Ye, Magnar Bjørås

## Abstract

We report a loss-of-function mutation in the TLDc domain of human *Oxidation Resistance 1* (*OXR1*) gene, resulting in early-onset epilepsy, developmental delay, cognitive disabilities, and cerebellar atrophy. Patient lymphoblasts show impaired cell survival, proliferation, and hypersensitivity to oxidative stress. These phenotypes are rescued by TLDc domain replacement. We generated patient derived induced pluripotent stem cells (iPSCs) revealing impaired neural differentiation along with dysregulation of genes essential for neurodevelopment. We identified that OXR1 influences histone arginine methylation by activating protein arginine methyltransferases (PRMTs), suggesting OXR1 dependent mechanisms regulating gene expression during neurodevelopment. We modeled the function of OXR1 in early human brain development using patient derived brain organoids revealing that OXR1 contributes to the spatial-temporal regulation of histone arginine methylation in specific brain regions. Our work provides new insights into pathological features and molecular underpinnings associated with OXR1 deficiency, highlighting the therapeutic potential of OXR1 in numerous neurodegenerative and neurodevelopmental disorders.

## Introduction

*Oxidation Resistance 1* (*OXR1*, HGNC: 15822, alias: *TLDC3*) is a highly conserved gene of TLDC domain containing family[1-6], first reported in *Escherichia coli* in 2000[7] and then confirmed in almost all eukaryotes[3]. OXR1 plays an essential role in oxidative stress resistance[3-20], but not directly in reactive oxygen species (ROS) scavenging[12] nor possessing catalase or superoxide dismutase activity[13]. It is involved in fundamental cellular and biological processes including DNA damage response[7, 8, 14], antioxidant pathways[14, 21], cell cycle[14, 15, 22], mitochondrial functions[8, 14, 16], glucose metabolism[23], S-nitrosylation[5], V-ATPase regulation[24], lysosomal functions[4], immune defense[25, 26], inflammatory response[18, 27], neuronal protection[3-5, 12, 17-20, 28], and life-span[13, 29]. It is associated to a wide range of diseases, such as amyotrophic lateral sclerosis(ALS) [3, 17-19], Parkinson disease[20], retinopathy[30, 31], and Lupus nephritis[27], pulmonary arterial hypertension[21], etc[21, 32, 33].

Due to the importance of ROS as signalling molecules in brain, the high metabolic rate, and a relatively low antioxidant capacity, the brain is assumed to be particularly susceptible to an imbalance in ROS levels[34]. Oxr1 depleted mice display progressive cerebellar ataxia, shortened lifespan and confers neuronal sensitivity to oxidative stress[5, 12]. Overexpression of Oxr1 in variant ALS mouse models was shown to delay onset of and improve motor neuron function and muscle pathology^14,16,17^. All Oxr1 isoforms are highly expressed in the central nervous system and disruption of the TLDc domain in Oxr1 leads to neurodegeneration[3]. In addition to the C-terminal TLDc domain, Oxr1 long isoforms also contain the evolutionarily conserved Tre2/Bub2/Cdc16 (TBC) domain and Lysin Motif (LysM)[3]. However, the functional roles of these domains remain elusive.

Very recently, it was reported that five patients from three families who carry four biallelic loss-of-function OXR1 variants were associated with cerebellar atrophy[4]. The impact of OXR1 deficiency was further assessed in a Drosophila model. Complete deletion of the *mustard* (*mtd*) gene, the fly homolog of human *OXR1* and *NCOA7* was lethal, which was rescued by the replacement of human *OXR1* cDNA (TLDC domain only). In addition, neuron specific knockout of *mtd* led to an accumulation of aberrant lysosomal structures, massive neuronal loss, and early death, but no significantly elevated oxidative stress was detected[4]. However, no human disease models are available to explore the pathological impact of OXR1 deficiency. The impact of OXR1 on cellular functions and molecular mechanisms in the human brain is largely unknown.

Here, we characterized a novel and deleterious OXR1 mutation in patients with severe neurological features. We produced patient-derived lymphoblasts, fibroblasts, iPSCs, and used the latter to generate monolayer neuronal cell cultures and various 3D brain organoid models (e.g., hippocampus, hypothalamus, midbrain and forbrain) to systematically analyze key aspects of the pathological features and molecular mechanisms associated with OXR1 depletion in brain. We found that the pathological phenotypes in patient lymphoblasts, such as impaired cell survival, proliferation, and hypersensitivity to oxidative stress, were rescued by the replacement of TLDc domain. In the patient derived iPSCs, impaired neural differentiation along with dysregulation of genes essential for neurodevelopment was identified. Further, we demonstrated that OXR1 influenced histone arginine methylation by activating protein arginine methyltransferases and revealed that OXR1 contributed to the spatial-temporal regulation of histone arginine methylation during human brain development.

## Materials and methods

### Whole exome analysis

Exonic sequences were enriched in the DNA sample of patient II-6 using SureSelect Human All Exon 50 Mb Kit (Agilent Technologies, Santa Clara, California, USA). Sequences were determined by HiSeq2000 (Illumina, San Diego, California, USA) as 100-bp paired-end runs. Data analysis including read alignment and variant calling was performed by DNAnexus software (Palo Alto, California, USA) using the default parameters with the human genome assembly hg19 (GRCh37) as reference. The study was performed with the approvals of the ethical committees of Hadassah Medical Center and the Ministry of Health (0393-17-HMO), and South-East Health Authorities of Norway (2014/658).

### Cell cultures

Immortalized lymphoblast cell (LB) lines were generated from patient II-4’s and healthy control’s peripheral blood mononuclear cells transformed with Epstein-Barr virus (EBV) as described by Bernacki SH et al[35], in the National Laboratory for the Genetics of Israeli Populations (NLGIP). The cells were cultured at 37°C with 5 % CO_2_ in growth medium: RPMI 1640 medium with GlutaMAX™ (Gibco, Cat.61870-010), supplemented with 20% fetal bovine serum (FBS), 100 IU/mL penicillin and 100 μg/mL streptomycin. Fibroblast cells and U2OS were cultured at 37°C with 5 % CO_2_ in growth medium: DMEM medium (Sigma-Aldrich, D6429) supplemented with 10% FBS. The trypan blue (0.4%, Invitrogen™) exclusion method is used to determine the number of viable cells present in a cell suspension. Cell counting was employed on the Countess^®^ automated cell counter (Invitrogen™).

### Plasmids and electroporation and cell sorting

The human cDNA of *OXR1A* ORF (NM_001198533) and the truncated variants of *OXR1A* lacking the TLDc domain, *OXR1A* ΔTLDc_1 and *OXR1A* ΔTLDc_2 consist of exon 1-16, 1-14, and 1-12, respectively, which were synthesized by GeneScript. They were inserted into plasmid pET28b and fused with the Hisx6 Tag between the Nde I and BamH I sites to obtain pET28b-*OXR1A* and pET28b-*OXR1A* ΔTLDc plasmids. All the plasmids were verified by sequencing. The plasmids of pET28b-*OXR1A, OXR1A*_ΔTLDc_1 and *OXR1A*_ΔTLDc_2 were transformed into *E. coli* and further used for purification of recombinant hOXR1 proteins by Ni-NTA and gel filtration columns.

Full length cDNA (NM_001198534) representing human *OXR1D*, synthesized by RT-PCR with mRNA from human HeLa cells, was clone in vector pEGFP-N1 and pEGFP-C1 with the GFP Tag between the Xho I and BamH I sites, to obtain pEGFP-N-*OXR1D* plasmids and pEGFP-C-*OXR1D*, respectively.

The pEGFP-C-*OXR1D* (pE-*OXR1D*) and The pEGFP-N2 (Vector) were transfected into patients’ LB (*OXR1*^ΔEx18^) by electroporation using a method modified from the patent (CN 103981218 A) ([Wenting Hou](http://www.pubfacts.com/author/Wenting+Hou), et al. Department of Biochemistry and Molecular Biology Peking University Health Science Center). LB-*OXR1*^ΔEx18^ in log phase growth were counted and harvested by centrifugation at 800 × g for 10 min at room temperature. After two washes with RPMI 1640 containing 10% FBS, the cells were resuspended in 37°C RPMI 1640 (with 10% FBS) at 1 × 10^7^ cells/mL. 250 μL of cell suspension and 15 μg of pEGFP-c-*OXR1D* or 12.5μg of pEGFP-N2 (Vector) were combined in a BTX 0.4-cm electroporation cuvette. Electroporation was done with ECM-830 (BTX^®^) at 210 V/pulse length=30ms/pulse=1. Electroporated cells were transferred to 2 mL of pre-warmed culture medium (with 10% FBS) at 37°C. The electroporated cells were allowed to grow in 5% CO_2_ at 37 °C for 48 hours(h) before cell sorting. Fluorescence-activated cell sorting (FACS)of EGFP-labeled LB were accomplished by FacsAria IIu high speed sorter at Flow Cytometry Core Facility, Oslo university hospital. The sorted cells were cultured at 37°C with 5 % CO_2_ in growth medium for two weeks. The expression level of *OXR1* was tested by real time qPCR.

### RNA isolation, cDNA sequence and gene expression analysis by Quantitative Real-Time Polymerase Chain Reaction

Total RNA was isolated from patient and control samples using the RNeasy Kit (Qiagen) following the manufacture’s protocol. The quality of purified RNA was estimated on an Epoch Microplate Spectrophotometer (Bio-Tek). Sequencing of *OXR1* cDNA from lymphocytes cDNA was prepared from fresh lymphocytes of patient II-4 and control using Maxima™ 1st strand cDNA synthesis kit (Thermo Scientific) and amplified by PCR using Phusion High-Fidelity PCR Master Mix (Thermo Scientific). The PCR products were separated using 2% (w/v) agarose gel electrophoresis and their sequence determined by Sanger sequencing. Two-steps qPCR method was used to determine mRNA levels. First, the cDNA was generated from total RNA samples using the High-capacity cDNA reverse transcription kit (Applied Biosystem). Real-time PCR reactions were prepared using Power SYBR Green Master mix including about 3 ng cDNA in each reaction, and qPCR was performed in the StepOnePlusTM Real-Time PCR System (Applied Biosystem) with the standard cycle conditions: 95 °C for 10 minutes(min); 40 cycles at 94 °C for 15 seconds(s) and 65 °C for 30 s. Samples were measured in triplicates. *ACTIN-ß* and *GAPDH* were used as internal standard, as indicated. Quantification was done using the comparative cycle threshold (CT) method of relative quantification (RQ), where RQ is the normalized signal level in a sample relative to the normalized signal level in the corresponding calibrator. The CT for the target gene was normalized to the CT for GAPDH in the same sample, giving ΔCT sample. The average ΔCT from cells’ cDNA was used as the ΔCT calibrator for the calculation of ΔΔCT. RQ is defined as 2−ΔΔCT, where ΔΔCT = ΔCT sample – ΔCT calibrator. Primers used to amplify target genes by qPCR are as follows: *HO-1* F (5’-AACTTTCAGAAGGGCCAGGT-3’), R (5’-CTGGGCTCTCCTTGTTGC-3’); *NRF2* F (5’-ACACGGTCCACAGCTCATC-3’), R (5’-TGTCAATCAAATCCATGTCCTG-3’); *p21* F (5’-GGCGGGCTGCATCCA -3’), R (5’-AGTGGTGTCTCGGTGACAAAGTC-3’); *OXR1* F (5’-TTATGGTACTGGAGAGACCTTTGTTTT-3’), R (5’-AAAACATATTATCTCCTGTCCACTTAAAGAC -3’); *FOS* F (5’-GGGGCAAGGTGGAACAGTTA-3’), R (5’-GTCTGTCTCCGCTTGGAGTG -3’); *c-JUN* F (5’-GTGCCGAAAAAGGAAGCTGG-3’), R (5’-CTGCGTTAGCATGAGTTGGC-3’); *DUSP1* F (5’-GGATACGAAGCGTTTTCGGC-3’), R (5’ -GGCCACCCTGATCGTAGAGT-3’); *CASP9* F (5’-TCTACGGCACAGATGGATGC -3’), R (5’-CATGGTCTTTCTGCTCCCCA -3’); *NANOG* F (5’-AATACCTCAGCCTCCAGCAGATG -3’), R (5’-TGCGTCACACCATTGCTATTCTTC -3’); *OCT4* F (5’-GTACTCCTCGGTCCCTTTCC -3’), R (5’-CAAAAACCCTGGCACAAACT -3’); *SOX2* F (5’-GGATAAGTACACGCTGCCCG -3’), R (5’-ATGTGCGCGTAACTGTCCAT -3’); *ACTIN* F (5’-GTTACAGGAAGTCCCTTGCCATCC -3’), R (5’-CACCTCCCCTGTGTGGACTTGGG -3’); *GAPDH* F (5’-CCACATCGCTCAGACACCAT -3’), R (5’-GCGCCCAATACGACCAAAT -3’).

### Western blot

Cell pellets were prepared using RIPA lysis buffer (150 mM NaCl, 50 mM Tris pH 7.5, 1mM EDTA, 0.1% SDS, 0.5% Sodium deoxycholate, and 1% Triton X-100, 1 mM DTT and 1x Protease Inhibitor Cocktail). The soluble protein extracts were separated by SDS-PAGE (4-15% mini-protein TGX precast gel, 12 or 15 well, BioRad) and transferred to a PVDF membrane (Trans-blot Turbo mini-Format 0.2 μM PVDF, cat.170-4156 by BIO-RAD). The membrane was blocked with 5% milk in PBS-T buffer (137mM NaCl, 2.7mM KCl, 4.3mM Na_2_PO_4_, 1.4mM KH_2_PO_4_, 0.1% Tween-20) for 4h at room temperature following incubation with the indicated primary antibodies, overnight at 4°C. For detection, membranes were incubated with secondary antibodies diluted in PBS-T buffer, except anti-GAPDH-HRP 1:5000 (Abcam Cat# ab9482, RRID:AB_307272). After washing 3 times with PBS-T buffer, the membrane was developed with ECL substrate (Super Signal West Fem to Maximum Sensitivity Substrate, by Thermo Scientific). The signals were detected by a BIO-RAD Imager and analyzed by Image Lab software. A list of antibodies used is provided in **Supplementary information**.

### Cell viability and apoptosis assay

The LB cells were plated in 6-well plates at 1×10^6^ cells/mL overnight. Then the cells were exposed to the indicated concentrations of H_2_O_2_ in fresh medium for 1 h and recovered for 24 h. After treatment, the cells were stained with propidium iodide (PI) and annexin V using Alexa Fluor 488 annexin V/Dead cell apoptosis kit (Invitrogen, Cat.V13245) according to the manufacture’s instruction. Analysis was performed via flow cytometry with BD Accuri C6 Sample software. At least 10,000 cells were analyzed for each sample.

### TUNEL staining

Apoptotic cell death was determined by In Situ cell death detection kit (Merck, Cat#11684795910) according to the manufacturer’s instructions. Briefly, cells were fixed with 4% paraformaldehyde and permeabilized with 0.1% Triton X-100 and 0.1% sodium citrate for 15 min. After washing, 100 μL of TUNEL reaction mixture was added and incubated for 1 h at 37°C. The cells were counterstained with 1 μg/mL DAPI (4′,6′-diamidino-2-phenylindole). The positive cells were imaged using an EVOS fluorescence microscope and analyzed using ImageJ software.

### Cellular general ROS measurement

The LBs in 6-well plates at 1×10^6^ cells/mL were incubated with 10 μM ROS probe CM-H2DCFDA (chloromethyl dichlorodihydrofluorescein diacetate, acetyl ester, Invitrogen, cat. 6827) in phosphate buffered saline (PBS) at 37°C for 30 min, washed once by PBS and recovered in phenol red free medium for 20 min. Next, the cells were exposed to 0.75 mM H_2_O_2_ in phenol red free medium for 1 h. Then, the cells were spin down and washed once with PBS, re-suspended in 0.5 mL cold PBS and kept on ice in dark. The fluorescence was measured using a BD Accuri™ C6 Cytometer.

### HPLC-MS/MS analysis of 8-oxo(dG) in genomic DNA

LBs were plated in 6-well plates at 1×10^6^ cells/mL overnight, and then exposed to the indicated concentrations of H_2_O_2_ in fresh medium and harvested after 1h, 4h and 6h treatment. Total DNA was isolated from LB using the DNeasy Kit (Qiagen) according to the manufacturer’s instructions. This assay consists of two independent experiments with technical triplicates (the first assay) or sextuplicates (the second assay).

For mass spectrometry analysis, genomic DNA was enzymatically hydrolyzed to deoxyribonucleosides by incubation for 40 min at 40 °C in a mixture of benzonase (Santa Cruz Biotechnology, sc-391121B), nuclease P1 from Penicillium citrinum (Sigma, N8630), and alkaline phosphatase from *E. coli* (Sigma Aldrich, P5931) in 10 mM ammonium acetate buffer pH 6.0 with 1 mM magnesium chloride. Proteineous contaminants were precipitated by addition of three volume equivalents of ice-cold acetonitrile and centrifugation at 16,000 g at 4°C for 40 min. Supernatants were collected and vacuum-dried at room temperature. The resulting residues were dissolved in water for HPLC-MS/MS quantification.

Chromatographic separation and mass spectrometry detection of 8-oxo(dG) was performed at room temperature using an Agilent 1290 Infinity II UHPLC system coupled with an Agilent 6495 Triple Quad (Agilent Technologies, Germany) and an Eclipse Plus C18 1.8 µm 150 × 2.1 mm i.d. column equipped with a guard column. The mobile phases used were: A (0.1 % formic acid in water), and B (0.1 % formic acid in methanol). Chromatographic separation started with 95% A and 5% B for 2.5 min; followed by linear gradients of: 0.5 min 5-13% B, 2.5 min 13-17% B, 1.5 min 45-65% B, 1.5 min 65-70% B, 1 min 70-5% B; 4 min re-equilibration with 5% B.

Chromatographic separation of unmodified nucleosides was performed using a Shimadzu Prominence LC-20AD HPLC system with an Ascentis Express C18 2.7 µm 150 × 2.1 mm i.d. column equipped with an Ascentis Express Cartridge Guard Column (Supelco Analytical, Bellefonte, PA, USA) with EXP Titanium Hybrid Ferrule (Optimize Technologies Inc.). Chromatographic separation was performed with isocratic flow of 25% methanol in water with 0.1 % formic acid, 0.16 ml/min at 40 °C. Online mass spectrometry detection of unmodified nucleosides was performed using an Applied Biosystems/MDS Sciex API5000 Triple quadrupole mass spectrometer (ABsciex, Toronto, Canada).

The deoxyribonucleosides were monitored by multiple reaction monitoring in positive electrospray ionization mode, using following mass transitions: 252.1→136.1 (dA), 228.1/112.1 (dC), 268.1/152.1 (dG), 243.1/127.0 (dT), 284.1/168.0 (8-oxodG).

### mtDNA mutation frequency measurement

Total DNA containing nuclear DNA and mtDNA were isolated from LB using the DNeasy Kit (Qiagen) according to the manufacturer’s instructions. Double stranded mtDNA mutation frequency was estimated as described previously[36]. Total DNA was digested with S1 nuclease (10U, Qiagen) for 15 min at 37°C and subsequently digested with TaqI restriction enzyme (100U, New England Biolabs) at 65°C for 1 h to remove single strands, damaged DNA and non-mutated DNA. The remaining mutated TaqI restriction fragments were quantified in a qPCR reaction. To ensure complete digestion of non-mutated TaqI restriction sites, TaqI was added to qPCR reaction mixture (100 U) and an additional step of 65°C for 15 min is included prior to the standard qPCR program. Mutation frequency is calculated as (2exp(CT_TaqI – CT_NT)*4)−1 per nt. The primer pairs for each gene were *12S* F (5’-AAACTGCTCGCCAGAACACT -3’) and R (5’-CATGGGCTACACCTTGACCT -3’); *D loop* F (5’-CCCGGAGCGAGGAGAGTAG -3’) and R (5’-CACCATCCTCCGTGAAATCAA -3’).

### Generation of human induced pluripotent stem cells (hiPSCs)

Fibroblasts from skin biopsy of patient II-2 and healthy control AG05836 (Coriell Institute) were cultured at 37°C with 5 % CO2 in growth medium: DMEM medium, supplemented with 10% fetal calf serum, 100 IU/mL penicillin and 100 μg/mL streptomycin. iPSCs were generated with CytoTune®-iPS 2.0 Sendai Reprogramming Kit (ThermoFisher Scientific, Cat#A16517) using a multiplicity of infection of 5:5:3 (KOS:c-Myc:Klf4). Individual colonies with typical morphology of human embryonic stem cells were manually picked around 3-4 weeks after infection. The iPSCs were continuously cultured and expanded on Geltrex coated 6-well plates with Essential 8 medium (ThermoFisher Scientific, Cat#A1517001). Human iPSC cell lines used in this manuscript: AG05836 clone 1, 10, 15, 27 as healthy controls and OXR1 clone 2, 10, and 11.

### Neural differentiation of hiPSCs

Differentiation of hiPSC towards neural lineage was performed as previously reported with modifications[37]. Briefly, iPSCs were pretreated with 50 μM Y-27632 (Stemcell Technologies) for 1 h. Then the cells were dissociated with Accutase (Stemcell Technologies) and plated onto low attachment V-type 96-well plate at 20 000 cells per well or AggreWell800 plate (Stemcell Technologies) to form aggregates. The culture was maintained in neural induction medium consisting of DMEM/F12 supplemented with 2% B27 supplement without vitamin A, 1% N2 supplement, 200 nM Noggin (R&D Systems) and 10 μM Y-27632 for 48 h. Formed aggregates were then transferred onto low attachment 6-well plates and continued culture for another 4 days. To induce neural rosette formation, the aggregates were transferred onto Geltrex coated 6-well plate and cultured for 4 days in neural induction medium supplemented with 200 ng/mL Noggin, 200 ng/mL DKK1 (R&D Systems), and 20 ng/mL bFGF (Life Technologies). Neural rosettes were manually picked and transferred onto low attachment 6-well plate in neural induction medium supplemented with 10 ng/mL bFGF and 10 ng/mL EGF to allow neuroepithelial sphere formation. Five days later, two procedures were performed to directly neural differentiation. First, the neural aggregates were placed onto Geltrex coated plates and treated with neural differentiation medium consisting of Neurobasal A medium supplemented with 10 ng/mL BDNF, 10 ng/mL GDNF, 200 μM ascorbic acid, 500 μM cyclic AMP. Second, the neural aggregates were dissociated into single cells with Accutase and replated onto poly-D-lysine/laminin coated 24 or 48-well plates for further differentiation with neural differentiation medium supplemented with 10 ng/mL BDNF, 10 ng/mL GDNF, 200 μM ascorbic acid, 500 μM cyclic AMP.

### Generation of cerebral organoids

Generation of cerebral organoids from iPSCs was performed based on an unguided approach previously described[38] with minor modifications to improve reproducibility, especially batch variations. Briefly, the neuroectodermal in the form of neuronal rosette were generated from multipotent iPSC aggregates (EBs) in suspension using minimalistic media, coupled with Matrigel embedding, and followed by the expansion of the elaborate architecture of the germinal zone of neural stem cells or progenitor cells with the differentiated neurons migration outwards containing the neural tissues of diverse identities or the potential to developing into multiple neural structures. Neuroepithelial buds developed further in a bioreactor. A major modification was using pipetting technic to remove all debris accumulated around the EBs by setting a multichannel pipette at 30 μL and gently pipetting up and down 5 times just before half medium replacement, which, we found, could largely improve EB’s quality. Another modification was to maintain the organoids in 96-well U-bottom ultra-low attachment plates until Matrigel embedding with half medium changing daily, aiming for individual organoid tracing and quality control.

### Generation of brain region-specific organoids

Generation of brain region-specific organoids from iPSCs was performed mainly based on a previous report[39] with modifications. The major modifications include: 1) culturing iPSCs with feeder free methods and generating EBs with Aggrewell™ 800 plates; 2) pattering starts since Day1; and 3) three-dimensional suspension culturing on an orbital shaker (Thermo fisher) with 120 rpm since Day2. Briefly, on Day 0, iPSC colonies were detached from Matrigel (Corning, cat. no. 354230) and disassociated by ReLeSR™ (Stemcell Technologies). After washed with fresh stem cell medium with 10 µM Y-27632, 10000 cells per microwell of Aggrewell™800 (3.0 ×10^6^ per well of 24-well in 2mL stem cell medium with 10 µM Y-27632) were seeded by centrifuging. From Day1, half of the medium was replaced following protocols diverge for the procedures to differentiate forebrain, midbrain and hypothalamus organoids developed by Qian *et al[40]*. The protocol of hippocampus organoids was developed by Fadi *et al[41]*. Briefly on Day2, around 30 organoids were transfer to each well of a 6-well plate placed on an orbital shaker rotating at 120 rpm in an incubator. For hippocampus organoids, on Days1-7, half medium was replaced with EB medium consisting of DMEM/F12 (Thermo Fisher Scientific), 1XGlutaMAX™ Supplement(Thermo Fisher Scientific), 20% (vol/vol) KnockOut™ Serum Replacement (Thermo Fisher Scientific), 1X Antibiotic-Antimycotic (Thermo Fisher Scientific), 1X 2-Mercaptoenhanol (Thermo Fisher Scientific), 1X Non-essential Amino Acids(NEAA, Thermo Fisher Scientific), 1 IU/mL Heparin (StemCell Technologies), 100 nM LDN193189 (StemCell Technologies), and 5 µM SB431542 (StemCell Technologies). On Days8-14, medium was exchanged to hippocampus patterning medium consisting of DMEM/F12, 1XGlutaMAX™ Supplement, 1XNEAA, 1X N2 (Thermo Fisher Scientific), 1 IU/mL Heparin, 2 µM CHIR99021 (StemCell Technologies), and 10 ng/mL Recombinant Human BMP-7(Peprotech). On Days15-44, hippocampus neurogenesis medium was applied, which consists of 1:1 DMEM/F12: Neurobasal Medium (Thermo Fisher Scientific), 1X GlutaMAX™ Supplement, 1X NEAA, 1X N2, 1 IU/mL Heparin, 1X B-27 Supplement Minus Vitamin A, and 2.5 mg/mL human insulin (Sigma-Aldrich). After Day 45, neurogenesis medium was exchanged with maturation medium consisting of 1:1 DMEM/F12: Neurobasal Medium, 1X GlutaMAX™ Supplement, 1XNEAA, 1X N2, 1 IU/mL Heparin, 1X B-27 Supplement Minus Vitamin A, 2.5 mg/mL human insulin, 20 ng/mL BDNF(Peprotech), 20 ng/mL recombinant Human GDNF (Peprotech), 200 µM Ascorbic Acid (StemCell Technologies), and 1µM Dibutyryl-cAMP (StemCell Technologies).

### Immunofluorescent staining

Immunofluorescent staining of neural cells were performed as previously described[42]. The cells were fixed with 4% paraformaldehyde (PFA) for 15 minutes, permeabilized with 0.1% Triton-X100 in PBS for 15 minutes and blocked with 5% bovine serum albumin(BSA), 5% normal goat serum, and 0.1% Triton X-100 in PBS at room temperature for 30 minutes, then incubated with primary antibodies overnight at 4 °C. After washing with 0.1% Tween-20/PBS, the samples were incubated with secondary antibodies for 1 h and counterstained with DAPI nuclear staining. Quantitative results were obtained with ImageJ software counting 10 random fields of each experiment.

Brain organoids were fixed with 4% PFA for 30 min at room temperature. Organoids were washed 3 times with PBS and immersed in a sucrose solution gradient, 10% for 30 min, 20% for 1 h, then in 30% sucrose solution overnight. Organoids were embedded in tissue freezing medium (Tissue-Tek® O.C.T. Compound) and sectioned with a cryostat (Thermo) at 16 μm thickness. For immunostaining, cryosection slides were laid at room temperate for at least 20 minutes, washed 3 times with PBS before being blocked with blocking medium consisting of 5% goat serum, 5% bovine serum albumin, 0.5 % Triton-X in PBS for 1 h followed by incubation with primary antibodies diluted in PBS buffer containing 0.5% goat serum, 0.5% BSA, 0.1% Tween 20 and 0.05% Triton-X at 4 °C overnight. For OXR1 staining, sections were then incubated at 90 °C in citrate antigen-retrieval buffer (pH 6.0) for 25 min and rinse once with PBS before blocking. After three washes in PBS containing 0.1% Tween 20, staining with secondary antibodies and DAPI was performed for 1 h at room temperature in the dark moisture chamber. After three washes in PBS containing 0.1% Tween 20 (10 min each), once quickly rinsing in PBS, and once quickly rinsing in pure water, the slides were mounted with ProLong™ Glass Antifade Mountant (Fisher Scientific, Waltham, Massachusetts). Cover glass thickness was 0.17 mm. The primary and secondary antibodies used, as well as their dilution are provided in **Supplementary information**.

### Analysis of immunolabelling

Images were captured on a Leica SP8 confocal microscope, using a 40x oil-immersion objective. Image analysis was performed in Fiji (ImageJ, NIH). Prior to analysis, Z-stacks were flattened with maximum or average z-projection. The same capture conditions and analysis settings were applied on both control and patient organoid sections. For quantification of progenitor cell proliferation and apoptosis, organoids were stained for KI67 and cleaved Caspase3(cCas3), respectively. Random fan-shaped regions from the apical surfaces of the ventricular zone (VZ) to the boundary between the outer subventricular zone (oSVZ)and cortical plate (CP) or between VZ and intermediate zone (IZ) in midbrain organoids were cropped for analyses. KI67^+^ nuclei and cCas3^+^ were counted and divided by the total number of nuclei stained by DAPI in the region respectively. To quantify the distribution of histone modifications in the developing cortical and midbrain organoids, radial structures were chosen randomly. From the pial surface to the basal surface, radial columns were cropped and evenly divided into 10-11 bins for analyses. The relative vertical position coordinates, y-coordinates on the image were recorded and normalized to the full thickness to measure their relative laminar positions in the differentiation and migration processes of midbrain progenitor cells or cortical laminar positions. The positions of each modification positive nuclei were separately marked using the Cell Counter plugin in Fuji(version 2.3.0). The positive cell proportion of the relative vertical positions in 10-11 evenly divided bins for each marker were calculated by (positive cell counts within each bin/total positive cell counts) and plotted in Prism (GraphPad 8.0). The mean intensity per cell were used to analysis the expression level of each marker.

### RNA sequencing and analysis

We conducted RNA-sequencing with three individual clones of each genotype at three stages of neuronal differentiation: iPSC (iPS_CTRL and iPS_*OXR1*^ΔEx18^), neural aggregate (NA) (NA_ CTRL and NA_ *OXR1*^ΔEx18^), and neuron (Neu) (Neu_CTRL and Neu_*OXR1*^ΔEx18^). in total, 18 samples were sequenced. Library construction and sequencing were performed through a commercially available service provided by BGI TECH SOLUTIONS (HONGKONG) CO. (Agreement No. F20FTSEUHT0295).

The quality control of fastq files was performed with FastQC v0.11.9[43]. Alignment to reference genome (GRCh38) was accomplished with hisat2 v2.1.0[44], while annotation and count matrix were computed with featureCounts v.2.0.0[45]. We accomplished downstream DEGs analysis in R v3.6.1 with DESeq2 v1.24.0[46]. Gene Set Enrichment Analysis(GSEA)[47] was performed to identify enriched pathways (using KEGG[48] and Reactome[49] databases) and gene ontology (GO) terms. Heatmaps were generated in R v3.6.1 with heatmap3 v1.1.7[50]. For the topologic biological network analysis, 1978 leading-edge genes determined by GSEA, which contributed the most to the top enriched GO terms in nervous system development, cell adhesion, extracellular matrix organization, neuron death, stress response, mitochondrial genome maintenance, glucose metabolism, lysosomal structure, and TFs from DEGs with |Log2FC| > 0.5 were recruited. Next, we constructed the protein-protein interaction (PPI) network for differential expression gene by searching STRING protein interaction database[51], with string score > 700 as the low limit value of confidence, FDR < 5%, and then performed network modules analysis by ReactomeFIViz[49]. A topologic biological network composed of the PPIs of 1539 DEGs clustering into 44 modules were established to help us infer the central transcriptome events. Each module consists of a set of genes that are both connected in the protein functional interaction network and highly-correlated in biological databases[52]. We run cytoHubba[53], TRRUST[54] and iRegulon(v1.3)[55] to identifiy hub genes, the drivers of a module and highly connected, and the essential upstream regulators. This network data were imported into and visualized by Cytoscape software v3.8.1[56].

### Proximity ligation assay

Proximity ligation assays (PLAs) were performed on fibroblast cells derived from the *OXR1*^ΔEx18^ patient and a healthy control. Cells were grown on glass coverslips, fixed with 4% PFA (added 1M glycine in the second wash step after fixation), and permeabilized with 0.1% Triton-X/1X PBS for 20min. The PLA was conducted according to the manufacturer’s instructions using the Duolink In Situ Red Starter Kit Mouse/Rabbit (Cat. No. DUO92101, SIGMA). After blocking for 30 min at 37°C, the cells were incubated with anti-OXR1 (1:200) and PRMT5 antibody (1:200) or anti-OXR1 and anti-PRMT1 antibody (1:200) overnight at 4°C. PLA Anti-Rabbit PLUS and Anti-Mouse MINUS probes containing the secondary antibodies conjugated with oligonucleotides, were added, and incubated at 37°C for ligation 30 min. Then the fluorescently labeled oligonucleotides resulting in red fluorescence signals were added to the reaction and incubated for amplification at 37°C for 100 min in the dark. The slides were mounted with Duolink *In Situ* Mounting Medium with DAPI (Cat. No. DUO82040, SIGMA). Microscopy was carried out using a Leica SP8 confocal microscope equipped with a 40X oil-immersion lens. The same input parameters were used throughout all experiments. Sample images were prepared in FUJI (ImageJ, NIH).

### Immunoprecipitation assays

The U2OS cell pellet was lysed with NETN buffer (20mM Tris-HCl pH 8.0, 100 mM NaCl, 0.5% NP-40, 0.5 mM DTT, 1:100 protease inhibitor cocktail, 400 U/mL). The lysate was pre-cleared by Dynabeads™ Protein A (Thermofisher Scientific, Cat#10001D) beads for 30 min. Add 1μg of purified rabbit anti IgG antibody (Sigma-Aldrich Cat# I5006, RRID:AB_1163659) or 3μg OXR1 antibody (Bethyl Cat# A302-036A, RRID:AB_1576565) to the 1mg pre-clear cell lysate separately. Incubate for 1h at 4ºC rotator. Add 20μL of the IP beads to the lysate and antibody mixture and incubate for overnight at 4°C with gentle agitation. Discard supernatant and wash the beads in 1x TBS buffer (50 mM Tris HCl pH 7.4, 150 mM NaCl, add 1 mM PMSF before used) 3 times. Denatured protein samples with SDS sample at 95°C for 10 mins. Load samples onto a 10% SDS-PAGE gel.

### Methylation assays with H4 N-terminal tail (aa 1-21) peptides

*In vitro* methylation assays were performed as previously described[1]. In a 30µL reaction, 7.35 nM PRMT5/MEP50 complex (Active motif, Cat#31356), 0.4 mM N-terminal H4 peptides, H4(1-21) WT and H4(1-21_R3K), with and without OXR1A or truncated OXR1A_ΔTLDC_1 or OXR1A_ΔTLDC_2, in a concentration gradient 1.83nM(+), 7.35nM(++) and 29.4nM(+++) respectively, were carried out in 50 mM HEPES (pH 8.0), 10 mM NaCl, and 1 mM DTT containing [3H] SAM (25:1 molar ratio of SAM(NEB, Cat#B9003S) to [methyl-3H]-SAM (PerkinElmer, Cat#NET155001MC). Reactions were incubated for 1 hr at 37°C, and then quenched with 0.5 µL of 100% trifluoroacetic acid. Each reaction was run in quadruplicate. Twenty-five microliters of each reaction were transferred to Merck Millipore MultiScreenHTS 96-Well Filter Plates with Negatively Charged Membrane (Cat#10245703) and air-dried for 30 min. The papers were subsequently washed in 50 mM NaHCO3 at pH 9.0 for 45 min, then air dried. Radioactivity was counted using 1450 MicroBeta Instrument (Wallac at Department of Immunology and Transfusion Medicine, Oslo University Hospital, Oslo, Norway).

### Quantification and statistical analysis

Each assay consists of at least two independent experiments with technical triplicates as described specifically. Data are presented as mean ± S.E.M., or mean ± S.D., as indicated in the figure legends. Organoid samples were randomly taken from the culture for experiments and analyses. Data analyses comparing control and OXR1 patient derived organoids were not performed blindly because the visual difference between the two groups is striking, and blinding was not possible to an informed researcher. Statistical analyses were performed using the Student’s *t* test in Excel or Graphpad Prism 8. A confidence level of 95% (\**p* < 0.05; ** *p* < 0.01, *** *p* < 0.001) was considered statistically significant.

## Results

### Characterization of OXR1 deficiency and description of clinical phenotypes

Three sisters whose parents were first cousins (**Fig. 1a**) presented delay or loss of developmental milestones after birth or at 8-10 months of age accompanied with focal myoclonic movement, hypoactivity, cognition problems, and frequent onset of epilepsy. Upon examination, patients showed non-communicative, generalized hypotonia, hyporeflexia/areflexia, ataxia and lack of spontaneous movements (**Table.1**). Electroencephalogram examination revealed generalized frank epileptic activity. Magnetic resonance imaging (MRI) disclosed atrophy particularly in vermis, cerebellum and thin corpus callosum (**Fig. 1b**). To identify the genetic lesion responsible for these clinical features, we first performed exome sequencing of patient II-6. Exome analyses revealed that the homozygous variant Chr8:107758095 G>C at the donor splice site located at exon 18 of the *OXR1* gene (Hg19 chr8:g.107758095G>C, XM_005250993.1:c.2552+5G>C) segregated with the disease (**Fig. 1c**). This mutation was not present in any of the nearly 60,000 exomes deposited by the Exome Aggregation Consortium, Cambridge, MA (URL: http://exac.broadinstitute.org, accessed August 2016). The human *OXR1* gene consists of 19 exons (ENSG00000164830, GRCh38), resulting in at least 4 major isoforms, OXR1A-D, with different tissue distributions. For instance, OXR1A and OXR1D are expressed in human brain[14], whereas most other cell types express isoform *B* and *D*. All isoforms contain the C-terminal TLDc domain. Sequencing cDNA of patient II-4 fibroblasts, the flanking *OXR1* exons 17-19 showed a shorter fragment (185 bp) compared to control cells (259 bp), indicative of homozygous skipping of exon 18 in *OXR1* (**Fig. 1d, Supplementary Fig.1a,b**), predicted to cause a frameshift leading to loss of the highly conserved C-terminal 69 amino acids of the TLDc domain of all OXR1 isoforms. OXR1 mRNA levels were similar in control and patient cells. Western blot of control cells showed a strong band corresponding to the OXR1B protein that was absent in patient cells (**Fig. 1e**), indicating that lack of exon 18 leads to protein instability. Therefore, it appears that the *OXR1* splicing mutation results in OXR1 depletion.

**Table 1.** Clinical features of OXR1 deficiency patients.

| Individual | II-3 | II-4 | II-6 |
| --- | --- | --- | --- |
| Inheritance pattern | Autosomal recessive inheritance |  |  |
| Pregnancy/At birth | Normal (Self-reported) |  |  |
| Age at symptom onset | Birth | 8 months | 10 months |
| Disease progress | No regression | Acute regression | Gradual regression |
| Clinical manifestations<br>Physical examination | Faltering growth |  |  |
|  | Mental retardation |  |  |
|  | Global neurodevelopmental delay, more pronounced in fine motor, speech and social skill |  |  |
|  | Epilepsy ( focal or myoclonic; refractory and frequent ) |  |  |
|  | Hypotonia, Hyporeflexia |  |  |
|  | Gait ataxia, Difficulties with balance |  |  |
| MRI | Vermis and cerebellar atrophy; Thin corpus callosum |  |  |
| EEG | Paroxysmal sharp slow waves |  |  |
| Important negative findings | No organomegaly |  |  |
|  | No malformation |  |  |
|  | Plasma lactate and urinary organic acids: normal |  |  |
|  | CSF analysis: normal |  |  |
|  | Vision was intact and fundoscopy: normal |  |  |

**Fig. 1:**
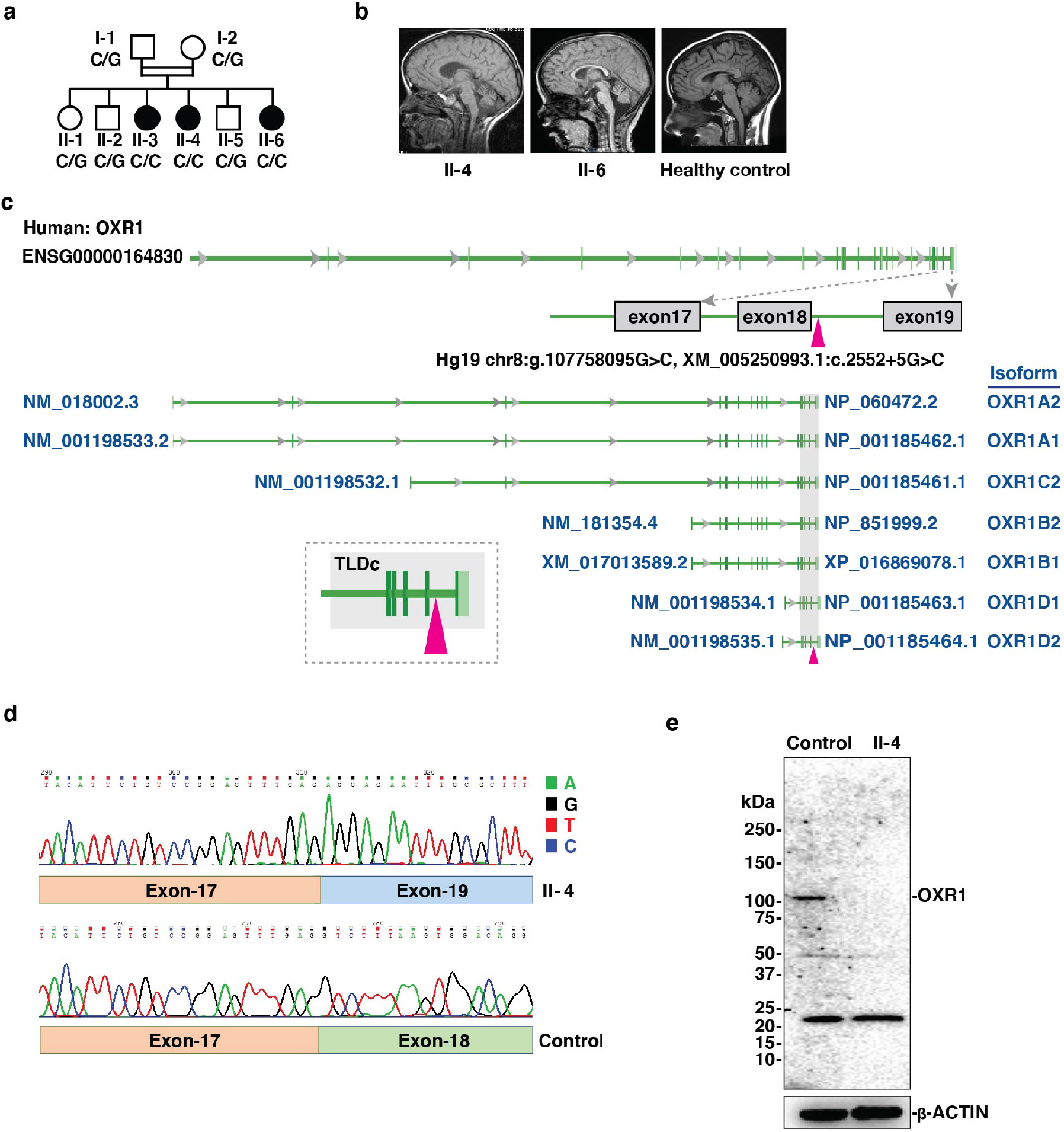
Donor splice site mutation in the *OXR1* gene leads to exon skipping and protein depletion. **a**, Family pedigree with genotype of Chr8:107758095 G>C mutation in the *OXR1* gene. **b**, Brain MRI. Mid-sagittal T1 weighted images showing marked cerebellar atrophy in patient II-4 at 6 years (left) and patient II-6 at 3 years and 4 months (middle), compared with the same view of a healthy control at 6-years (right). **c**, Genomic structure of human *OXR1* and the mutation site. The *OXR1* gene is located on chromosome 8 and consists of 19 exons, reference to ENSG000001164830 (Hg19). Triangle in magenta, the mutation in the donor splicing site of exon 18 within TLDc domain (*OXR1*^ΔEx18^), affecting all identified human *OXR1* isoforms. **d**, cDNA sequence around the *OXR1* mutation site of lymphoblasts from patient II-4 and healthy control. **e**, Western blot analysis of OXR1 level in lymphoblasts from patient II-4 and healthy control. β-actin was used as loading control.

### OXR1 deficiency compromises cell growth, survival, and response to oxidative stress in patients derived lymphoblasts

To explore the functional impact of OXR1 deficiency in patients, we established lymphoblast cell lines (LB) from patient (LB*-OXR1*^ΔEx18^) and from one unaffected sibling as control (LB-CTRL). The LB*-OXR1*^ΔEx18^ showed reduced proliferation (**Fig. 2a**), higher sensitivity to H_2_O_2_ with increased apoptosis and necrosis (**Fig. 2b, Supplementary Fig. 2a**). H_2_O_2_ treatment induced higher ROS levels in LB-*OXR1*^ΔEx18^ compared to control (**Supplementary Fig. 2b**). To determine the levels of oxidative DNA damage, we measured accumulation of 8-oxoguanine (8-oxoG) in genomic DNA[57]. Basal levels of 8-oxoG were similar in patient and controls, but increased 8-oxoG levels were observed in LB*-OXR1*^ΔEx18^ upon H_2_O_2_ exposure (**Fig. 2c**). Furthermore, we determined the rate of mitochondrial DNA (mtDNA) mutagenesis[36], showing elevated mutation frequencies in the D-loop and 12S regions of mtDNA in LB*-OXR1*^ΔEx18^ (**Supplementary Fig. 2c**). We have previously shown that OXR1 upregulates the expression of antioxidant genes via the p21 signaling pathway in Hela cells[14]. In this study, we observed a reduction of both p21 and heme oxygenase (HO-1) proteins in LB*-OXR1*^ΔEx18^ (**Fig. 2d**) and upregulation of stress response genes, including *DUSP1*, and the transcription factors (TF) *FOS* and *JUN* at the mRNA level (**Fig. 2e**). ROS induced activation of caspases is pivotal in apoptosis. Caspase-9 mRNA levels (**Fig. 2f**), as well as the activated forms of Caspase-9 (35 and 37 kDa) were significantly increased in LB*-OXR1*^ΔEx18^ (**Fig. 2g**). Taken together, these results demonstrate that OXR1 deficiency reduces cell proliferation and increases intracellular oxidative stress, oxidative DNA damage and apoptosis.

**Fig. 2:**
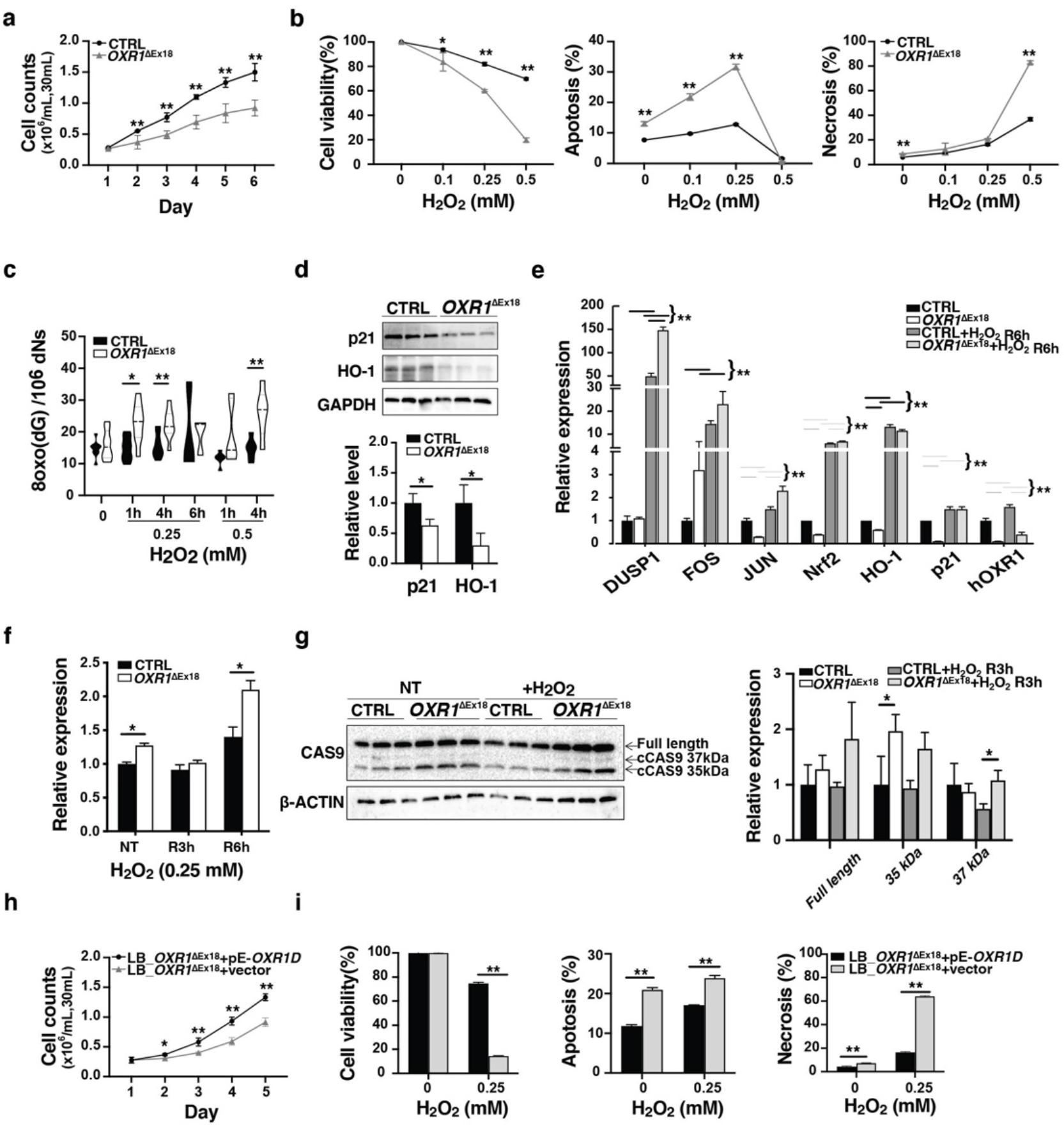
OXR1 regulates cell proliferation, anti-oxidation response and apoptosis. **a**, Proliferation capacity of patient lymphoblasts (*OXR1*^ΔEx18^) as compared with control cells (CTRL). n=3 independent experiments, n=6 samples for each genotype in each experiment. **b**, Patient (*OXR1*^ΔEx18^) and control (CTRL) lymphoblasts were treated with indicated concentrations of hydrogen peroxide (H_2_O_2_) for 1 h and harvested after 24 h recovery, populations of surviving (left), apoptotic (middle) and necrotic (right) cells determined by flow cytometry with FITC-Annexin V/PI double staining. n=3 independent samples. **c**, 8-oxoG levels in genomic DNA detected by mass spectrometry after H_2_O_2_ treatment in patient (*OXR1*^ΔEx18^) and control (CTRL) lymphoblasts. n=3 independent samples in the first assay (0.25 mM), and n=6 independent samples in the second assay. **d**, Quantification of p21 and HO-1 protein expression measured by western blot analysis under regular culturing conditions of patient (*OXR1*^ΔEx18^) and control (CTRL) lymphoblasts. n=3 independent samples. **e**, Quantitative real-time PCR analysis of oxidative stress response genes in patient (*OXR1*^ΔEx18^) and control (CTRL) lymphoblasts exposed to H2O2 and recovery for 6 hours (R6h). n=3 independent samples. **f**, qPCR analysis of Caspase-9 (*CAS9*) expression in patient (*OXR1*^ΔEx18^) and control (CTRL) lymphoblasts following 0.25 mM H_2_O_2_ treatment and recovery for 3 hours (R3h) or 6 hours (R6h). n=3 independent samples. **g**, Western blot analysis and quantification of CAS9 expression and cleaved CAS9 (cCAS9) in patient (*OXR1*^ΔEx18^) and control (CTRL) lymphoblasts exposed to 0.25 mM H_2_O_2_ and recovery for 3 hours (R3h) or non-treated (NT). β-ACTIN used as loading control, n=3 independent samples. **h**, Proliferation capacity of patient lymphoblasts with stable expression of OXR1D (LB*-OXR1*^ΔEx18^+pE-OXR1D) compared to patient lymphoblasts carrying only vector (LB-*OXR1*^ΔEx18^+vector), n=6 independent samples. **i**, Analyses of cell viability, apoptosis and necrosis patient lymphoblast samples stably expressing OXR1D (LB*-OXR1*^ΔEx18^+pE-OXR1D) or empty vector (LB-*OXR1*^ΔEx18^+vector) upon H_2_O_2_ treatment.n=3 independent samples. All Data are shown as mean±SD. \**p* < 0.05, \*\**p* < 0.01, Student’s *t* test.

The TLDc domain is essential for OXR1’s anti-oxidation function[3]. OXR1D is predominantly composed of the TLDc domain and is expressed in all types of tissues in human. To validate whether the observed cellular phenotypes were due to the mutation in the TLDc domain, we performed rescue experiments using LB*-OXR1*^ΔEx18^ cells stably expressing OXR1D-GFP. Strikingly, expression of OXR1D-GFP was sufficient to increase the proliferation capacity of LB*-OXR1*^ΔEx18^ (**Fig. 2h**) and reduced apoptosis and necrosis to levels comparable to control LBs (**Fig. 2i**). These results suggest that the splicing site mutation in TLDc domain leads to depletion of OXR1 isoforms with loss-of-function.

### Loss of OXR1 causes morphological and developmental defects in neuronal cells

To overcome the challenge due to the lack of access of patient brain tissues for understanding the pathological mechanisms, we generated iPSCs from fibroblasts of patient II-4 and a healthy control (AG05836). Loss of exon18 was conserved, and the OXR1 protein was absent in the patient iPSCs (**Supplementary Fig. 3a,b**). All control and *OXR1*^ΔEx18^ clones showed normal karyotypes, similar expression levels of pluripotency markers and *in vitro* confirmation of pluripotency via directed differentiation to the three germ layers (**Supplementary Fig. 3c-f, Supplementary video1**). However, further differentiation of iPSCs into neurons revealed irregularities in the morphology of OXR1^ΔEx18^ neural aggregates (NAs) (**Supplementary Fig. 3g,h**). Moreover, OXR1-depletion reduced neurite growth (**Fig. 3a**), as well as the number of neurons, concomitant with reduced expression of neuronal markers (**Supplementary Fig. 3i, j**). Similar to patient LBs (**Fig. 2b**), OXR1-depleted neuronal cells displayed enhanced apoptosis (**Supplementary Fig. 3k**). Notably, *OXR1* mRNA and protein levels increased during neural differentiation in control cells (**Supplementary Fig. 3l**). Taken together, these results demonstrate an essential role of OXR1 in neural differentiation.

**Fig. 3:**
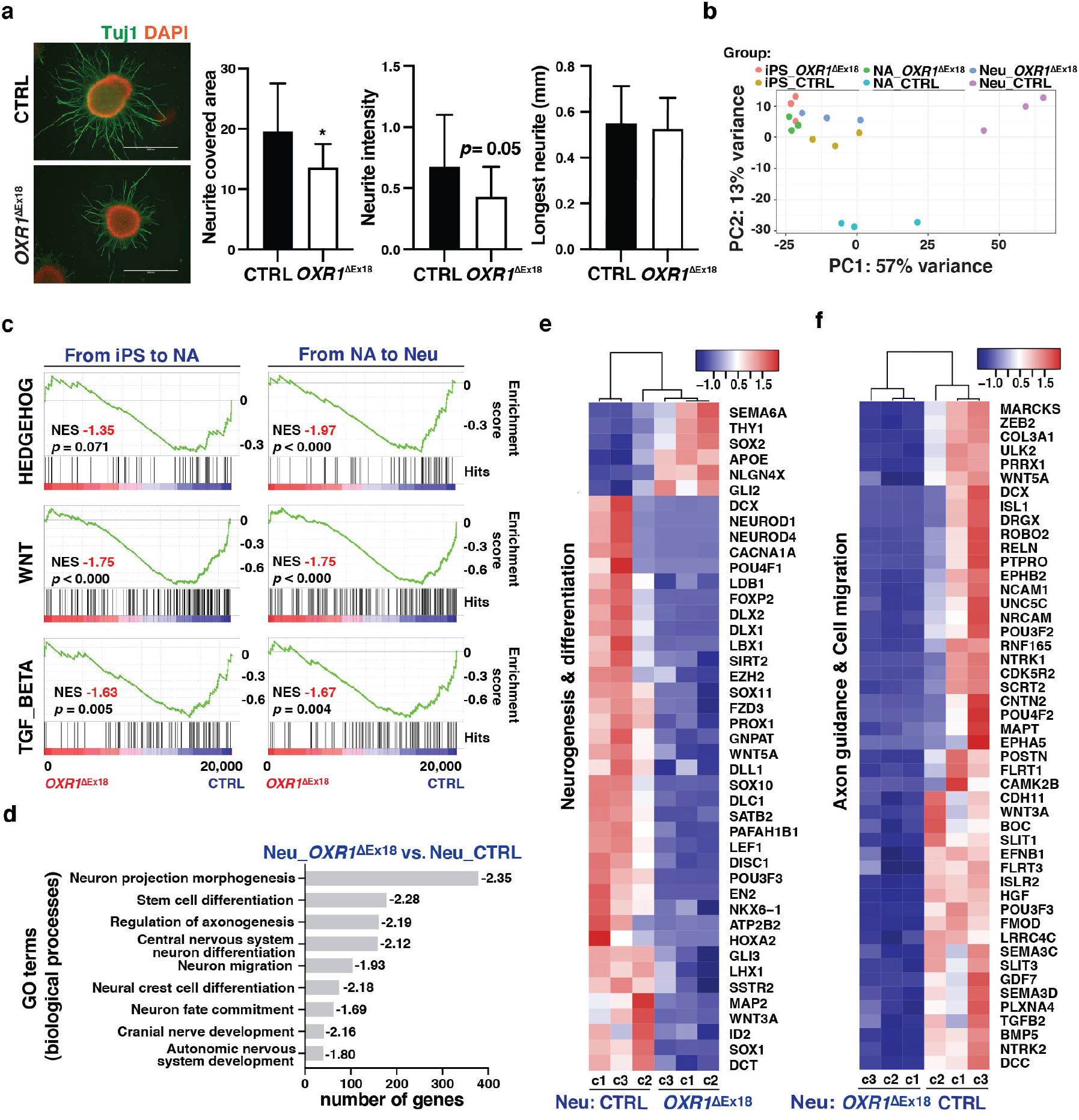
OXR1 is required for Planar neural differentiation. **a**, Immunocytochemical analyses of neurospheres from patient (*OXR1*^ΔEx18^) and healthy control (CTRL). Neurite outgrowth is visualized by antibody against neuron-specific class III beta-tubulin (Tuj1). Bar diagrams show quantification of the neurite (Tuj1^+^) covered area, the fluorescence intensity of covered area, and the longest neurite length by average of 5 longest neurites of each neurosphere (n = 15 independent neurospheres for each genotype). **b**, Principal components analysis (PCA) showing the clustering of samples based on principal component 1 and 2 (PC1 and PC2). All samples are three different clones of each genotype derived from the monolayer neuronal differentiation experiment. Induced pluripotent stem cells (iPSC) were collected at day0, neural aggregates (NA) at day2 (48h after initiation of differentiation) and neurons (Neu) at day20. **c**, Pathway enrichment analysis showing the core pathways involved in neural development by Gene Set Enrichment Analysis (GSEA, normalized enrichment score, |NES| ≥ 1; nominal *p* value, (*p*) < 0.05; false discovery rate (FDR) ≤ 0.25). **d**, Gene Ontology (GO) analysis by GSEA. The most underrepresented GO terms of biological processes in *OXR1*^ΔEx18^ neurons are relevant to neural development. **e**, Heatmap showing z-score normalized read counts of the top ranked leading-edge genes in neurogenesis and differentiation that were differentially expressed in Neu_*OXR1*^ΔEx18^. **f**, Heatmap showing z-score normalized read counts of the top ranked leading-edge genes in axon genesis and cell migration that were differentially expressed in Neu_*OXR1*^ΔEx18^.

### OXR1 impacts transcription during neuronal differentiation

To elucidate OXR1-dependent regulation of transcriptional networks during neural differentiation, transcriptome analysis were performed at three distinct stages during neuronal differentiation (iPSCs, NAs and neurons) of control and *OXR1*^ΔEx18^ iPSCs (**Supplementary Fig. 3h,i-u**). Transcriptomes of three clones of control iPSCs, NAs and neurons showed distinct clustering, suggesting a well-defined, stage-specific neural differentiation process (**Fig. 3b**). In contrast, no prominent clustering was observed between the three groups of *OXR1*^ΔEx18^ cells. From iPSCs to neurons via NAs, lack of OXR1 lead to progressive failure in neuron-fate commitment (**Supplementary Fig. 3n**), suggesting impaired neuronal differentiation trajectories upon OXR1 depletion. Pathway enrichment by Gene Set Enrichment Analysis (GSEA) showed that the core pathways involved in neural development, such as Hedgehog, WNT, and TGF-beta, were activated in control, but not in *OXR1*^ΔEx18^ (**Fig. 3c**). Most differentially expressed genes (DEGs) due to OXR1 depletion were identified at the neuron stage (**Supplementary Fig. 3n**). Gene Ontology by GSEA revealed underrepresentation of biological processes required for the nervous system development, including hindbrain development (**Fig. 3d, Supplementary Fig. 3p-q**). Some overrepresented biological processes such as synaptic signaling and neurotransmitter transport were also identified (**Supplementary Fig. 3o**), indicating a potential role of OXR1 in synaptic regulation. Genes essential for neurogenesis and neurodifferentiation as well as axon guidance and cell migration were significantly down-regulated in Neu*-OXR1*^ΔEx18^ (**Fig. 3e-f**). In addition, we found that several pro-apoptotic genes were upregulated while anti-apoptotic genes were down-regulated in Neu-*OXR1*^ΔEx18^ (**Supplementary Fig. 3r**), potentially contributing to the increased death of neurons **(Supplementary Fig. 3k)**. Several oxidative stress response genes were upregulated (**Supplementary Fig. 3s**), supporting the oxidative distress in OXR1-depleted neurons, as observed in LB*-OXR1*^ΔEx18^. According to a topologic biological network analysis of the dysregulated transcriptome of Neu*-OXR1*^ΔEx18^, we found that OXR1 orchestrated the expression of many key upstream regulators crucial for regulating transcription and substantial pathways for neural differentiation (**Supplementary Fig. 3t**). Further biological module clustering directly revealed OXR1-depletion impaired axon guidance at multiple levels, from axon genesis to neurite extension and synapse formation[58]. Among them, the dysregulated Reelin pathway, cell adhesion molecules (ie. *NRCAM*), Class-3 semaphorin, and Slit/Robo pathways were highlighted, which correlated to the formation of a thinner corpus callosum[59],[60, 61].

### OXR1 modulates PRMT-mediated histone modifications

Next, we explored molecular mechanisms underlying OXR1-dependent regulation of gene expression. Previous mass spectrometry analysis revealed the interaction of overexpressed OXR1-GFP in U2OS cells with protein arginine methyltransferase 5 (PRMT5)[1], an epigenetic regulator of gene expression. Here, we confirmed the interaction of endogenous OXR1 with PRMT5 as well as PRMT1 in U2OS cells by immunoprecipitation (IP) (**Supplementary Fig. 4a**), and in human fibroblasts (control vs *OXR1*^ΔEx18^) by an *in-situ* proximity ligation assay (**Fig. 4a**). PRMTs can activate or repress gene expression by catalyzing histone (H) arginine (R) methylations on the N-terminus of histones H3 and H4, e.g., H3R2, H3R17, and H4R3[62]. Crosstalk between histone arginine methylation and lysine acetylation/methylation has also been reported[63]. We examined histone modifications during differentiation of iPSCs to neural stem cells (NSCs) using an antibody-based screening and revealed increased histone methylations on H3R2/R17, H4R3, H4K4/K9/K27 in NSCs of the control (**Fig. 4b**). In contrast, OXR1-depleted NSCs showed significantly impaired histone arginine/lysine methylations, e.g., H3R2me2a, H3R17me2s/2a, H4R3me2s/2a, H3K4me3, H3K9me3, and H3K27me3 as compared to control cells. However, we did not detect any changes in the expression of PRMTs in either *OXR1*^ΔEx18^ iPSCs or NSCs as compared to control cells, except PRMT7 that showed significantly lower levels in *OXR1*^ΔEx18^ NSCs (**Supplementary Fig. 4b**). Thus, our results suggest that OXR1 may mainly affect the catalytic activity of PRMT-induced epigenetic histone modifications during neurogenesis.

**Fig. 4:**
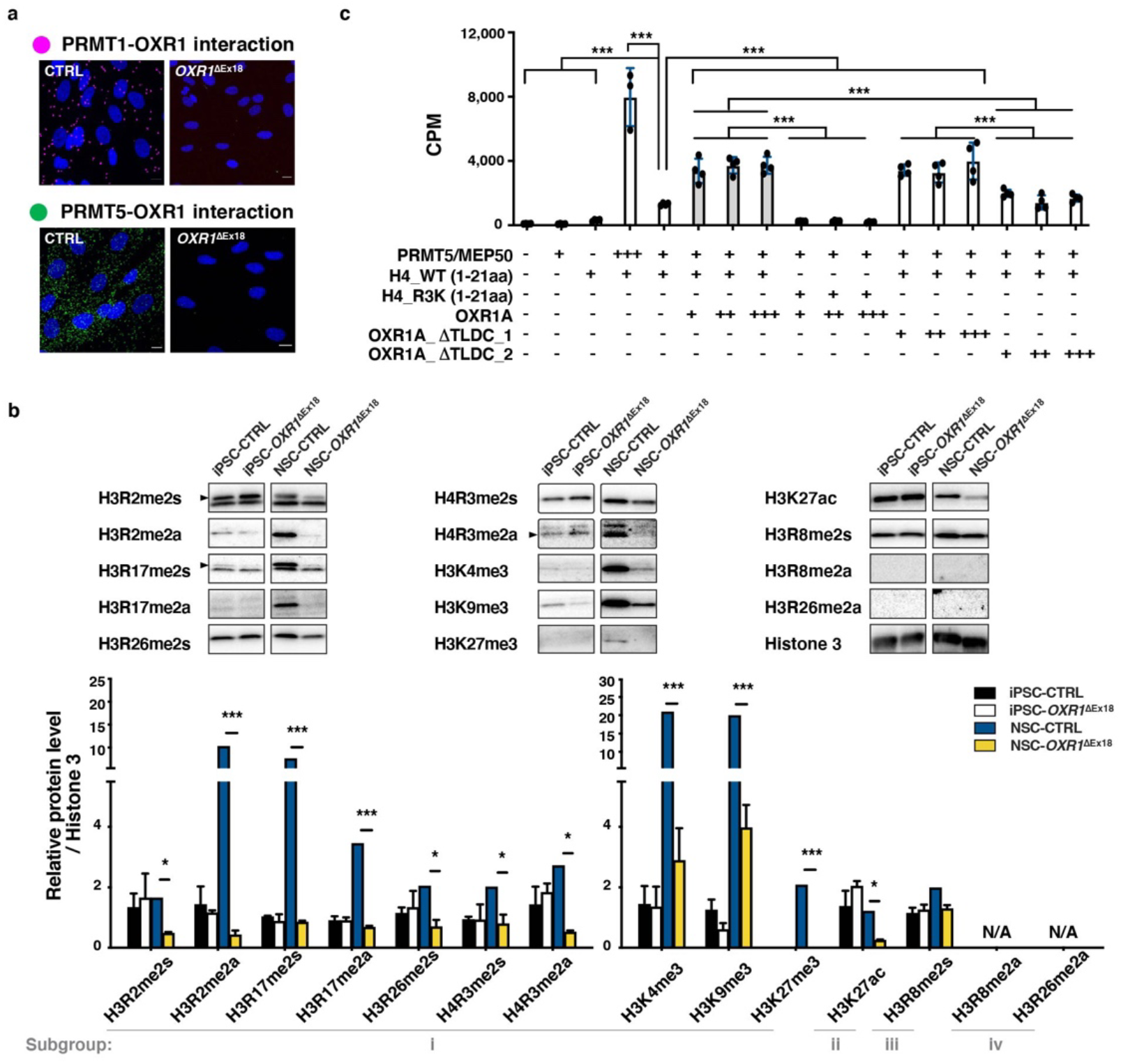
OXR1 modulates PRMT-induced epigenetic histone modifications during neurogenesis. **a**, Proximity ligation assay showing the interaction of OXR1 with PRMT1 (magenta spots) and PRMT5 (green spots) in patient fibroblasts (*OXR1*^ΔEx18^) and control cells (CTRL). Nuclei are indicated by DAPI staining. **b**, The Western blot analysis showing histone arginine modifications in induced pluripotent stem cells (iPSC) and neural stem cells (NSC). The arginine modification marks include H3R2me2s, H3R2me2a, H3R17me2a, H4R3me2a, H4R3me2s, H3R17me2s, H3R26me2s, and lysine marks H3K4me3, H3K27me3, and H3K9me3; Histone 3 was used as loading control. **c**, *In vitro* methyltransferase assay showing a direct role of OXR1 in regulating the enzymatic activity of PRMT5. N-terminal H4 peptides (1-21aa) and the mutant ones H4-R3K (1-21aa) were used as substrates. Different molar ratio of PRMT5/MEP50 protein complex (calculated as tetramer) to the full length OXR1A or truncated ones, such as 4:1 (+), 1:1(++), and 1:4(+++), was applied in each reaction. The enzymatic activity of PRMT5 was shown by the CPM (Count per minute) value of ^3^H-Methyl incorporation. All Data are shown as mean±SD, \**p* < 0.05, \*\*\**p* < 0.001, Student’s *t* test.

To gain more insight into how OXR1 regulates histone marks, we integrated our RNA-sequencing data with published ChIP-sequencing datasets using the standard quality control available from http://cistrome.org/db. We found that genes enriched with the repressive mark H4R3me2s overlapped with 71% of the down-regulated genes during neural differentiation in control cells (1319 out of 1869 genes), of which 63% of the genes were differentially expressed (up-regulated) in *OXR1*^ΔEx18^ cells (802 out of 1266 genes). Furthermore, genes enriched with H3K4me3, H3K9me3, H3K27me3, and H3K27ac overlapped with 16%-27% of the down- or up-regulated DEGs in *OXR1*^ΔEx18^ cells. These data may suggest a role of OXR1 in regulation of histone modifications, particularly H4R3 di-methylation.

To further explore a functional role of OXR1 in PRMT-catalyzed methylation of H4R3, we assessed the activity of the PRMT5/MEP50 methyltransferase on the Histone 4 N-terminal tail (aa1–21) in a methyltransferase assay *in vitro* by adding to the reactions full-length hOXR1A or mutant proteins with partial or complete deletion of TLDc (**Fig. 4c**). Interestingly, full-length hOXR1A and the mutants with partially truncated TLDc (OXR1A_ΔTLDc_1) stimulated PRMT5-catalyzed H4 methylation by 3-fold. In contrast, the OXR1A mutant lacking TLDc (OXR1A_ΔTLDc_2) failed to stimulate PRMT5 activity, suggesting the TLDc domain of OXR1 is essential for regulation of PRMT-catalyzed H4R3 methylation.

### Impaired development of OXR1 deficient human brain organoids

To unravel pathological processes at molecular, cellular, and structural levels, we modeled OXR1 deficiency in the developing human brain organoids. We established cerebral organoids[38], and various brain region-specific organoids[39, 41]derived from patient iPSCs. Cerebral organoids often contain heterogeneous tissues resembling various brain regions, which relies on the intrinsic differentiation signaling and self-organization capacities of iPSCs (**Fig. 5a**). Significantly, *OXR1*^ΔEx18^ cerebral organoids were smaller than the healthy controls (CTRL) with slower growth rate and suboptimal morphology of neuroepithelial buds (**Fig. 5a**). After about 30 days, the cortical regions in healthy controls (**Supplementary Fig. 5a**) displayed dense and radially organized ventricular subventricular zone (SVZ) like structures harboring the neural progenitor cells (NPCs). The neuronal layer composed of TBR1^+^ and CTIP2^+^ deep-layer neurons was highly reminiscent of the formation of cortical plate (CP). New-born neurons continued to migrate radially outward, became mature (MAP2^+^) and obtained rudimentary separations during the lamination process (**Fig. 5b**). However, in *OXR1*^ΔEx18^ organoids, the two layers still intermingled at day75, indicating either delayed neurogenesis or impaired neuron migration, or a combination. In addition, we observed a thick Reelin-positive (RELN^+^) layer along the basal surface reminiscent of marginal zone (MZ) with long and extended RELN^+^ plexus at inner layer in control organoids at day75 (**Fig. 5c**). It is known that RELN expressed at the marginal zone is essential for correct cortical and cerebellar neuronal lamination[64], and RELN localization along dendrites suggests a role in synaptic remodelling and neuronal maturation[65]. However, thinner MZ with reduced RELN deposition, shorten axons and lack of RELN^+^ plexus was observed in patient iPSC-derived organoids. These results suggest that OXR1 deficiency leads to delayed neuronal layering and in human cerebra.

**Fig. 5:**
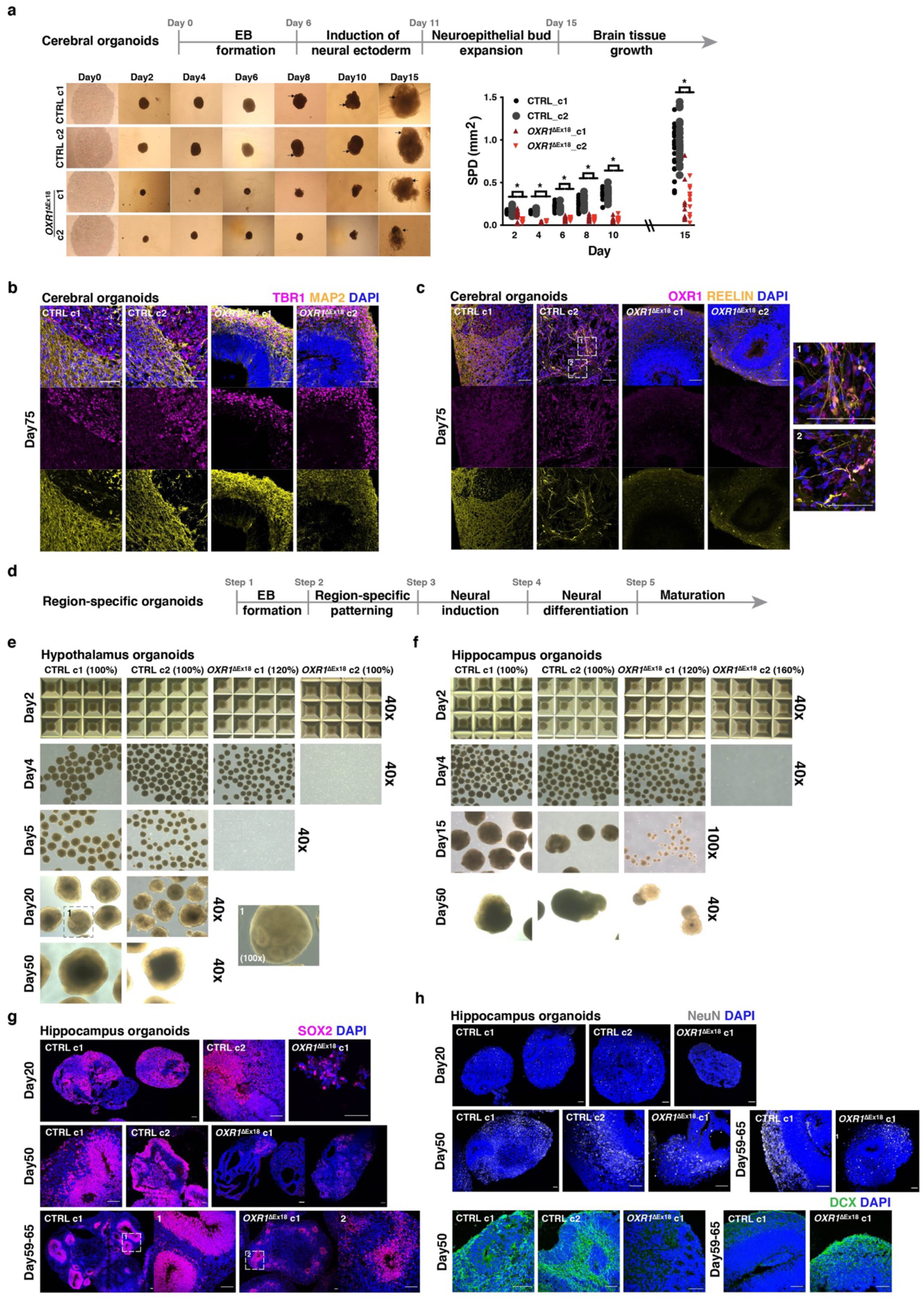
OXR1 deficiency severely impairs brain development in distinct human brain organoid models. **a**, The progression of cerebral organoid development is illustrated (upper). Representative images (bright field, 40x) show the progression of cerebral organoids derived from each healthy control (CTRL) and patient (*OXR1*^ΔEx18^) iPSC clones from day 2 to day 15 (lower left). Black arrows point the expanding neuroepithelial formation in CTRL clones from day 8 and the poorly developed neuroepithelia buds observed in the *OXR1*^ΔEx18^ cerebral organoids at day 15. The body size of cerebral organoids (lower right) is quantified by summing the products of bi-dimensional measurements (SPD). SPD = Longest diameter (LD) Δ longest diameter perpendicular to the LD. n=15-20 organoids of each clone, * *p* < 0.05. **b**, Immunohistochemical analyses of cerebral organoids at day 75 using the antibody specific for MAP2 (the marker of mature neurons, yellow) or TBR1(the marker of deep layer neurons, magenta). **c**, Immunohistochemical analyses of cerebral organoids at day 75 using antibodies specific for REELIN (yellow) and OXR1 (magenta), respectively. **d**, A schematic illustration showing the guided progression of region-specific brain organoids. **e**, Representative images (bright field, 40x) show the progression of hypothalamus derived from healthy control (CTRL) and patient (*OXR1*^Δ*Ex18*^) iPSCs. The starting cell number of each clone is indicated (100% refer to 10,000 iPSC cells per microwell of Aggrewell-800 plate on day 0). After hypothalamus patterning, the spheroids of *OXR1*^ΔEx18^ clone 1 (c1) and *OXR1*^ΔEx18^ clone 2 (c2) dissipated by day 5. **f**, Representative images (bright field, 40x) show the progression of hippocampus organoids. After hippocampus patterning, the spheroids of *OXR1*^ΔEx18^ c2 dissipated by day 5. **g**, Representative confocal images showing the progression of SOX2^+^ neural progenitor cells (magenta) during the development of hippocampus organoids. **h**, Representative confocal images showing the progression of NeuN^+^ mature neurons (grey) and DCX^+^ immature neurons (green) during the development of hippocampus organoids. In all confocal images, the scale bar is 60 µm and the nuclei were visualized using DAPI (blue).

*OXR1*^ΔEx18^ patients showed a global neurodevelopmental delay charactered by the dysfunctions of cognition, fine motor skills, learning and language, suggesting impaired functions not only in cerebellum, but also in cortical (i.e, prefrontal) areas and deep structures (i.e, hippocampus, basal ganglia). To assess the importance of OXR1 in distinct brain regions, we guided neural progenitor cells (NPCs) to form brain region-specific identities by supplement of highly tailored external patterning factors at the early stage (**Fig. 5d-h, 6a-d**) [39, 66, 67]. From control iPSCs, we successfully generated hypothalamus, hippocampus, forebrain (cortical) and midbrain organoids with large VZ-like structures containing the region-specific NPCs. Molecular markers for cell identities were analyzed by IHC of different region-specific organoids. OXR1 is highly expressed in both POMC^+^ peptidergic neurons and OTP^+^ hypothalamic neurons of hypothalamus organoids at day 50 (**Supplementary Fig. 5b**). Strikingly, OXR1-deficient clones from patient iPSCs lost the ability to form hypothalamus organoids (**Fig. 5e**). In addition, iPSC*-OXR1*^ΔEx18^ generated hippocampal organoids from only one clone (**Fig. 5f**) with a reduced size of SOX2-positive progenitor zone, a less organized layer of DCX-positive newborn neurons, and fewer NeuN-positive mature neurons (**Fig. 5g-h**). Taken together, these results suggest a crucial role of OXR1 in early hypothalamus and hippocampal development.

**Fig. 6:**
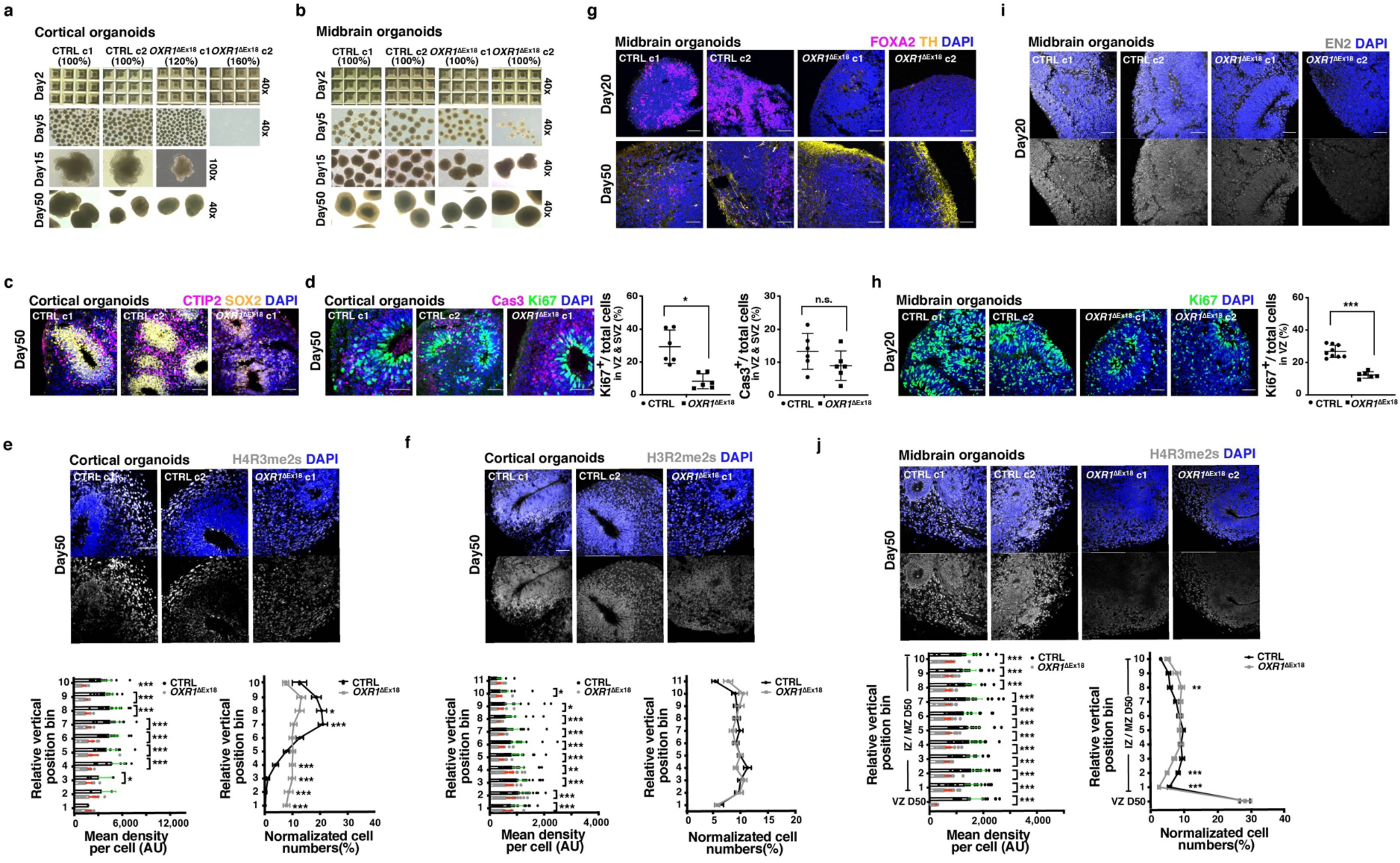
OXR1 impacts cortical and midbrain development by shaping histone arginine methylations. **a-b**, Representative bright-field images (40x) show the progression of cortical and midbrain organoids, derived from healthy control (CTRL) or and patient (*OXR1*^Δ*Ex18*^) iPSCs. The starting cell number of each clone is indicated (100% refer to 10,000 iPSC cells per microwell of Aggrewell-800 plate on day 0). After cortical patterning, the spheroids of *OXR1*^ΔEx18^ clone 2 (c2) usually dissipated by day 5. After midbrain patterning, *OXR1*^ΔEx18^ c2 showed less midbrain organoids generated at day 5. **c**, Immunohistochemical analyses showing the CTIP2^+^ deep layer neurons (magenta) formed above SOX2-enriched progenitor zones (yellow) of cortical organoids at day 50. **d**, Immunohistochemical analyses showing the Ki67^+^ proliferating cells (green) and cCas3^+^ apoptotic cells (magenta) at VZ/SVZ progenitor zones of cortical organoids at day 50. The percentage of Ki67^+^ or cCas3^+^ cells in VZ/SVZ is quantified as shown in the bar diagram. n = 6 organoids. VZ, ventricular zone; SVZ, subventricular zone. **e**, Immunohistochemical analyses showing the distribution of H4R3me2s^+^ cells in cortical organoids at day 50. The average expression of H4R3me2s in each cell is quantified in each vertical position bin (lower left). The normalized abundance of H4R3me2s^+^ cells in each vertical position bin is calculated as the number of H4R3me2s^+^cells in a bin / the total number of H4R3me2s^+^ cells. CTRL, n=17; OXR1^ΔEx18^, n=14 from two independent batches. **f**, Immunohistochemical analyses showing the distribution of H3R2me2s^+^ cells of cortical organoids at day 50. The average expression of H4R3me2s of each cell is quantified in each vertical position bin (lower left). The normalized abundance of H3R2me2s+ cells in each vertical position bin is calculated as in (**e**). CTRL, n = 37; *OXR1*^*ΔEx18*^, n=38 from two independent batches. **g**, Immunohistochemical analyses of midbrain organoids at day 20 and day 50 using antibodies specific for FOXA2 (magenta, the marker of midbrain floor plate/progenitor) and TH (yellow, the marker of dopaminergic neurons). **h**, Immunohistochemical analyses showing Ki67^+^ proliferating cells (green) in the VZ of midbrain organoids at day 20. The percentage of Ki67^+^ cells in progenitor zones (VZ) is quantified. CTRL, n=9 organoids; *OXR1*^*ΔEx18*^, n=6 organoids. **i**, Immunohistochemical analyses showing transient expression of EN2 (the marker of mid-hindbrain boundary) in midbrain organoids at day 20. **j**, Immunohistochemical analyses showing the spatial-temporal distribution of H4R3me2s in midbrain organoids at day 50. The average expression of H4R3me2s in each cell is quantified in each vertical position bin (lower left). The normalized abundance of H4R3me2s^+^ cells in each bin is calculated as in (**e**). CTRL, n = 23; *OXR1*^*ΔEx18*^, n=23 from two independent batches. **d-f, h-j**, Values represent mean ± SEM, \**p <* 0.05, \*\**p <* 0.01, *\*\*\*p <* 0.001, *Student’s t* test. In all confocal images, the scale bar is 60 µm and the nuclei were visualized using DAPI (blue).

### OXR1 impacts cortical and midbrain development by shaping histone arginine methylations during neurodifferentiation

We successfully generated cortical (forebrain) and midbrain organoids from control and *OXR1*^ΔEx18^ iPSCs (**Fig. 6a, b**). Control cortical organoids contained the large and radially organized VZ/SVZ like structures marked by SOX2 (**Fig. 6c**), where cortical NPCs underwent active cell division (marked by Ki-67, in **Fig. 6d**). The newly generated layer of CTIP2^+^ early born neurons extended the formation of CP resembling human early corticogenesis (**Fig. 6c**). Interestingly, we identified a specific pattern for the histone arginine methylation mark H4R3me2s, with low levels in VZ/SVZ like structures but gradually increasing in developing neurons outside the progenitor zone following the radial migration (**Fig. 6e**). This finding suggests a regulatory role of H4R3me2s during neurogenesis and the establishment of the cortical plate. In contrast, this pattern of H4R3me2s was lacking in *OXR1*^ΔEx18^ cortical organoids showing an overall reduction in the level of H4R3me2s as compared to control organoids (**Fig. 6e**). In addition, *OXR1*^ΔEx18^ cortical organoids showed a thin and poorly organized VZ/SVZ with markedly reduced number of Ki67^+^ proliferating NPCs, and a barely developed CTIP2^+^ cortical plate (**Fig. 6c, d**). H3R2me2s, another target of PRMT5, displayed an even distribution pattern across the normal laminar cortical structures of control and *OXR1*^ΔEx18^. However, *OXR1*^ΔEx18^ cortical organoids showed an overall reduction in H3R2me2s as compared with control organoids (**Fig. 6j**). These results suggest that histone arginine methylations play roles in human early corticogenesis, and that OXR1 deficiency impairs histone arginine methylation leading to abnormal cortical neurogenesis and CP formation.

Midbrain neurogenesis starts at VZ of the midbrain floor plate (mFP) where midbrain progenitors are differentiated to various types of tyrosine hydroxylase (TH^+^) midbrain dopaminergic (mDA) cells (**Supplementary Fig. 6a**). We guided midbrain NPCs towards the identity of ventral tegmental area to generate *substantia nigra pars compacta* (SNc) mDA neurons via the Shh-FOXA1/2-SOX6 pathway by using SHH, CHIR99021 and purmorphamine as patterning factors[39, 58, 68-74]. As expected, FOXA2^+^ progenitors were broadly detected at day 20, migrated through the intermediate zone (IZ) and became TH^+^ mDA when reaching mantle zone (MZ) at day 50 (**Fig. 6g**). In *OXR1*^ΔEx18^ midbrain organoids, FOXA2^+^ progenitors were sparsely detected at day 20 (**Fig. 6g**) with fewer Ki67^+^ proliferating NPCs (**Fig. 6h**), although the structural organization of VZ remained largely intact. In contrast, we observed an abnormal increase of TH^+^ mDA generated at day 50 (**Fig. 6g**), suggesting dysregulation of midbrain NPCs specification derived from the activation of FOXA2. It is known that upregulation of EN2 by FGF8 in the midbrain-hindbrain boundary is essential for further cerebellar development[75-77]. Similarly, EN2 was induced in VZ in control organoids, but absent in *OXR1*^ΔEx18^ (**Fig. 6i**), suggesting a potential mechanism underlying the cerebellar defects in patients. Notably, the regulation of H4R3me2s in the normal midbrain development was distinct from forebrain by showing a high level in the VZ progenitor zone **(Fig. 6j)**. Nevertheless, the level of H4R3me2s was dramatically decreased in *OXR1*^ΔEx18^ organoids. However, we did not detect any significant changes in H3R2me2a or H3R2me2s *OXR1*^ΔEx18^ midbrain organoids as compared to control organoids (**Supplementary Fig. 6b,c**). Interestingly, we found both the distribution pattern and the expression level of histone lysine mark H3K4me3 were severely dysregulated in *OXR1*^ΔEx18^ midbrain organoids (**Supplementary Fig. 6d**). Taken together, these results suggest that OXR1 modulates distinct histone marks during early region-specific brain development, accounting for the clinical phenotypes in *OXR1*^ΔEx18^ patients.

## Discussion

Here we report a *de novo* mutation within the critical TLDc domain of *OXR1* leading to exon skipping, protein depletion (*OXR1*^ΔEx18^) and a severe early-onset syndrome characterized by epilepsy, global developmental delay, cognitive disabilities, and cerebellar atrophy. Using 2D neural differentiation and various 3D brain organoid models derived from patient iPSCs, we identified specific neurodevelopmental defects due to OXR1 depletion, including impairment of NPC proliferation, fate-specification, neurogenesis, neurite outgrowth, and neuronal migration. We revealed a role of OXR1 in epigenetic regulation of early human brain development by modulating PRMT-catalyzed histone arginine methylation.

Searching the Genome-wide association study (GWAS) database showed that genetic variants of *OXR1* are associated with several common neurological disorders, e.g. amyotrophic lateral sclerosis (ALS, HGVST65/375), Alzheimer’s disease (HGVST51), Parkinson disease (PD, HGVST6/83) or ischemic stroke (HGVST14). Further, search in published databases showed that DEGs of *OXR1*^ΔEx18^ patient derived neuronal cells overlap 39%, 41% and 31% with disease associated genes or high-confident risk genes of cerebellar atrophy, autism spectrum disorder and schizophrenia, respectively[78-80] (**Supplementary Fig. 7**), supporting a role of OXR1 in protection against a broader spectrum of neurodevelopmental disorders. Three children from the same family have been identified carrying a stop-gain mutation very near the start of *OXR1D* (non-TLDc region) with dyslexia or specific language impairment[81]. Furthermore, four loss-of-function variants in *OXR1* of five individuals from three independent families have been reported[4]. Each of the five individuals carry bi-allelic variants in *OXR1* showing similar clinical manifestations as the *OXR1*^ΔEx18^ patients reported in this work.

The highly conserved TLDc domain is common for six different proteins (e.g., TBC1D24 and NCOA7), which all appears to display neuroprotective roles according to recent studies[3, 82]. A mutation in TBC1D24 was the first genetic cause of human epilepsy found[83]. Later, more than 50 mutations in TBC1D24 have been associated with various types of drug-resistant epilepsy, intellectual disability, and DOORS (<u>d</u>eafness, <u>o</u>nychodystrophy, <u>o</u>steodystrophy, mental <u>r</u>etardation, and <u>s</u>eizures) syndrome, of which at least 12 mutations locate in TLDc domain[84, 85]. In this work, advanced human cell culture systems modeling OXR1 depletion provide new insight into the molecular and cellular pathology of rare diseases associated with inherited mutations of the TLDc containing family. Previous studies show that loss of OXR1 in various animal models and cell lines leads to neurodegeneration[3-5, 12, 17-20, 28]. Most of these studies demonstrate that the TLDc domain is essential for protection of neurons against oxidative and nitrosative stress-induced cell death, e.g in mice cerebellum[3, 5, 12, 19, 23]. Other studies suggest that OXR1 could regulate glucose metabolism in mice cerebellum[23] or lysosomal structure and function in *Drosophila* neurons[4]. Here, human *OXR1*^ΔEx18^ patient derived cells showed elevated oxidative stress and increased cell apoptosis in both lymphoblasts and 2D differentiated neurons, and dysregulation of transcriptional networks indicating impaired mitochondrial function, glycolysis and autophagy (**Supplementary Fig. 3t**). According to BrainSpan Atlas of the developing human brain[86], OXR1 expression increase largely from the early second trimester, reaching a plateau at two years of age until brain development is completed. Redox homeostasis is crucial for brain development, termed ‘oxidative eustress’[87]. We identified several key modulators of redox homeostasis that were significantly affected in Neu*-OXR1*^ΔEx18^, including H_2_O_2_ scavengers, key redox-sensitive targets and redox hubs (**Supplementary Fig. 3s,t**), suggesting an important role of OXR1 not only in maintaining the balance between oxidative eustress and distress, but also in the proper redox signaling required for neuron development[87-90]. There is a strong increase in oxidative eustress at PCW 14–16[91] coinciding with cortical organoids around day 50. This developmental stage may therefore be particularly dependent on OXR1 mediated redox signaling. Taking the advantages of iPSC derived brain organoids in recapitulating the complexity of early corticogenesis, we revealed that OXR1 deficiency resulted in delay of specific cellular programs, including NPC proliferation, CP formation, cortical layering, and axon/dendrites growth. Impaired region-specific NPC proliferation and neurogenesis have been closely linked to primary microcephaly and vermis/cerebellar hypoplasia[92, 93]. Errors in the movement or placement of incoming neurons may therefore have consequences for cortical networks, leading to seizures and cognitive disability[93].

Arginine methylation, a widespread post-translational modification catalyzed by PRMTs, plays key roles in epigenetic regulation and signal transduction influencing gene transcription, RNA metabolism and DNA damage response. Dysregulation of histone arginine methylations has been linked to pathomechanisms of several diseases, including neurodegeneration (i.e., PRMT1-ALS, PRTM5-PD), and muscular disorders [17, 62, 94, 95]. Studies in mouse, other animal models and mouse derived NSC/NPC cultures have demonstrated roles of PRTM1/3/4/5/7/8 on the development of glial or neuronal lineages[62]. PRMT5 depletion in mouse brain leads to depletion of the external granular layer of the cerebellum, reduced thickness of the cortex with lower number of neuronal cells in CP and SOX2/Ki67-positive proliferating NPCs in VZ/SVZ, and death within 14 days after birth[96]. However, to our knowledge there are no reports on PRMTs roles in human brain development. In 2D derived NSC*-OXR1*^ΔEx18^, we showed that multiple histone arginine methylations were dysregulated, but levels of their relevant PRMTs were not affected. We found strong overlap between H4R3me2s associated genes and DEGs in OXR1 depleted neurons. Moreover, OXR stimulated the enzyme activity of PRTM5 of H4R3me2s *in vitro*. We propose that OXR1 is a coactivator of several PRMTs regulating histone arginine methylations ensuring a proper transcriptome for neurodevelopment. Recent studies have demonstrated that brain organoids possess comparable transcriptomic profiles to fetal brain at certain stages, as well as the similar regulation of epigenetic programs. For example, the transcriptome and ChIPseq of H3K4me3 and H3K27ac of cortical organoids at around day 50 mapped most consistently to isogenic cortical brain tissues at PCW16[79, 80]. Further, we monitored the spaciotemporal changes of histone marks during brain development using brain organoids, especially focusing on epigenetic regulation of cortical or midbrain NPCs at the early stage. Notably, we revealed that histone arginine modifications are active in the early stage of human cortical and midbrain development in a region-specific way. Loss of OXR1 severely interfered with histone modifications including both arginine and lysine marks, which may lead to the dysregulated transcriptome underlying impaired neurogenesis and neurodegeneration in *OXR1*^ΔEx18^ patients.

The roles of arginine methylation have been more challenging to decipher than other PTMs because of the broad specificity of PRMTs, which target both histone and non-histone proteins[94]. Notably, many pathways affected by OXR1 deficiency are modulated by arginine methylation of non-histone proteins by PRMTs^*[62, 95, 97, 98]*^, including WNT, BMP, p53, and RNA metabolism and splicing (Module 5,17,18, **Supplementary Fig. 3t**). Thus, how OXR1 affect non-histone arginine PTMs during neural differentiation merits further investigations.

## Acknowledgements

We thank the family of the described patients for participating in our studies. Funding was provided by the Research Council of Norway, South-East Norway Regional Health Authority. The Proteomics and Metabolomics Core Facility PROMEC is funded by NTNU and the Central Norway Regional Health Authority, and is a member of the National Network of Advanced Proteomics Infrastructure (NAPI), which is funded by the RCN INFRASTRUKTUR-program (295910). The authors wish to acknowledge the help and resources provided by the National Core Facility for Human Pluripotent Stem Cells, Norwegian Centre for Stem Cell Research, Oslo University Hospital, in particular the use of the virus facility and confocal microscope suite.

## Author Contributions

XL, WW, MB, ND, MY and OE were responsible for concept and design of the study and involved in all aspects of the project. OE collect clinical information and generated lymphoblast and fibroblast cell lines from OXR1 patients and their healthy siblings. XL and MY performed experiments with LBs. AK and GS provided the 8-oxoG measurements in LBs genomic DNA. WW and GJS generated iPSC cell lines. WW and XL performed 2D neuronal differentiation experiments. MP and XL carried out RNA-seq analysis. XL and WW performed brain organoid experiments. F.J., H.S., G-l.M. contributed to brain organoid studies. MB and XL performed the methyltransferase assay. AARI, RS, and LE performed clinical data collection. MS, MAA, BAL, SE, and OE conducted clinical examination. XL, WW, JY, GJS, MMLS, BVL, SOB, WT, JEH, OE, AL, SE and MB wrote and revised manuscript. All authors reviewed the manuscript and approved the final version.

