## Supplement Figures for "Loss-of-function mutation in human *Oxidation Resistance gene 1* disrupts the spatial-temporal regulation of histone arginine methylation in early brain development"

**a**

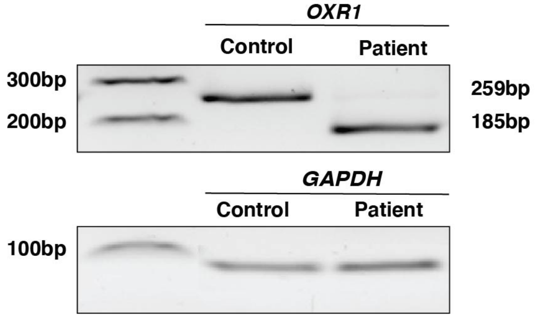


**b**

<u>GTC TTT AAG TGG ACA GGA GAT AAT ATG TTT TTT ATC AAA GGA GAC ATG GAT TCA CTA</u>
<u>GCT TTC GGT GGT GGA GG</u>A GGA GAA TTT GCG CTT TGG CTT GAT GGA GAT CTC TAC CAT
GGA AGA AGC CAT TCT TGT AAA ACG TTT GGG AAT CGT ACA CTT TCT AAG AAG GAA GAT
TTC TTT ATC CAA GAT ATT GAA ATC TGG GCT TTT GAA TAA

**Supplementary Figure 1. Donor splice site mutation in the *OXR1* gene causes TLDc domain depletion. a,** PCR amplified cDNA fragment spanning exon 17-19. The shorter (185bp) product in the patient sample indicates the deletion of exon 18. **b**, The absence of exon 18 results in an open reading frame shift and consequently a premature stop codon “TGA” in exon 19. The underlined part is exon 18, while the rest is exon 19.

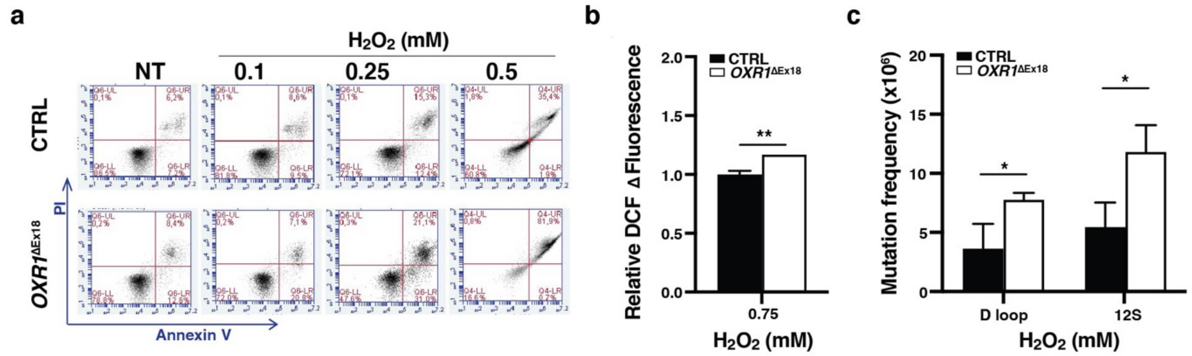


**Supplementary Figure 2. Loss of OXR1 increases cell death, general ROS level and mitochondrial DNA mutation frequency in lymphoblasts (LB). a,** Representative results of FITC-Annexin V/PI double staining by flow cytometry (FCM). **b**, Elevated amounts of ROS generated in patient LB cells (*OXR1*$^{\Delta \text{Ex18}}$) after $H_2O_2$ exposure, as compared with the changes in control (CTRL). Intracellular ROS level was measured by DCFDA staining via FCM analysis. **c**, Increased mitochondrial DNA mutation frequency in D-loop and 12S regions in *OXR1*$^{\Delta \text{Ex18}}$ under normal culturing condition. **b-c**, n=3 independent experiments, * $p < 0.05$, ** $p < 0.01$, *t*-test.

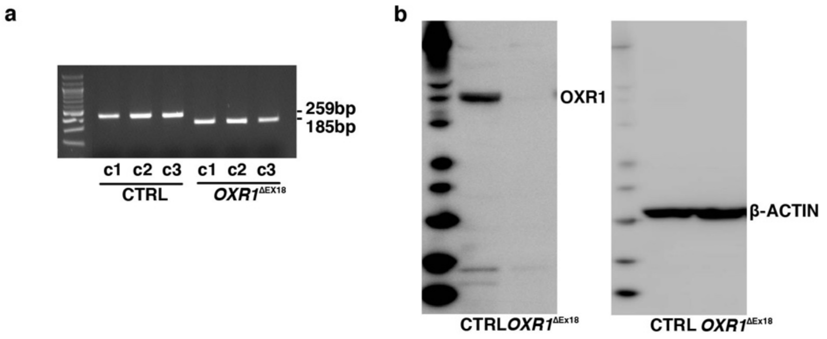


c

**Karyotype** — **Median ratios**

| Sample | Gender | CV% | Threshold ; Threshold Y | 01C | 02C | 03C | 04C | 05C | 06C | 07C | 08C | 09C | 10C | 11C | 12C | 13C | 14C | 15C | 16C | 17C | 18C | 19C | 20C | 20q1 | 21C | 22C | Auto | X | Y |
|---|---|---|---|---|---|---|---|---|---|---|---|---|---|---|---|---|---|---|---|---|---|---|---|---|---|---|---|---|---|
| AG-F_1 | Female | 5.92 | 0.88-1.12 ; 0.82-1.18 | 0.97 | 1.06 | 0.98 | 0.97 | 0.97 | 0.96 | 1.01 | 1.03 | 1.05 | 1.08 | 1.07 | 1.04 | 1.02 | 0.94 | 1.08 | 1.02 | 1.07 | 1.00 | 1.08 | 1.04 | 1.05 | 1.05 | 1.05 | 0.94 | 1.23 | 0.73 |
| AG-F_2 | Female | 6.67 | 0.87-1.13 ; 0.80-1.20 | 1.02 | 1.04 | 0.94 | 1.02 | 0.97 | 0.98 | 1.05 | 1.00 | 1.03 | 1.05 | 1.03 | 1.04 | 1.02 | 1.08 | 1.04 | 0.99 | 0.99 | 0.90 | 1.05 | 0.98 | 1.00 | 1.05 | 1.00 | 1.00 | 1.21 | 0.81 |
| Oxr-F_1 | Female | 4.99 | 0.90-1.10 ; 0.85-1.15 | 0.98 | 0.92 | 0.95 | 1.00 | 0.97 | 0.94 | 0.99 | 0.99 | 0.97 | 1.03 | 0.98 | 0.99 | 0.99 | 1.00 | 1.06 | 0.98 | 0.98 | 0.96 | 1.02 | 0.98 | 0.98 | 1.00 | 0.95 | 1.07 | 1.21 | 0.76 |
| Oxr-F_2 | Female | 5.60 | 0.89-1.11 ; 0.83-1.17 | 0.98 | 1.02 | 1.01 | 0.96 | 0.99 | 1.03 | 0.99 | 1.01 | 0.99 | 1.00 | 0.99 | 1.07 | 1.01 | 1.01 | 1.08 | 1.02 | 0.95 | 0.96 | 1.02 | 1.00 | 1.09 | 1.00 | 0.99 | 1.00 | 1.24 | 0.77 |
| AG-c10_1 | Female | 6.35 | 0.87-1.13 ; 0.81-1.19 | 1.01 | 1.04 | 1.00 | 0.98 | 0.96 | 0.97 | 1.01 | 1.01 | 1.05 | 1.06 | 1.02 | 0.98 | 0.98 | 1.04 | 1.06 | 1.00 | 1.01 | 1.00 | 1.04 | 1.00 | 0.95 | 0.98 | 0.94 | 1.01 | 1.16 | 0.77 |
| AG-c10_2 | Female | 6.75 | 0.87-1.13 ; 0.80-1.20 | 1.00 | 0.99 | 1.02 | 0.96 | 0.96 | 0.98 | 0.97 | 0.94 | 1.06 | 1.05 | 1.06 | 1.01 | 1.02 | 1.04 | 1.06 | 0.95 | 0.95 | 1.00 | 1.02 | 1.01 | 0.93 | 0.96 | 0.92 | 1.01 | 1.15 | 0.71 |
| AG-c27_1 | Female | 6.44 | 0.87-1.13 ; 0.81-1.19 | 0.99 | 0.99 | 0.99 | 0.97 | 0.96 | 0.99 | 0.99 | 0.98 | 0.99 | 1.04 | 1.02 | 0.97 | 0.98 | 0.93 | 1.03 | 0.96 | 0.98 | 0.93 | 1.06 | 0.98 | 1.00 | 0.97 | 0.91 | 1.07 | 1.13 | 0.77 |
| AG-c27_2 | Female | 5.68 | 0.89-1.11 ; 0.83-1.17 | 1.07 | 1.02 | 1.00 | 1.06 | 1.02 | 1.00 | 0.98 | 1.01 | 1.09 | 1.08 | 1.04 | 1.07 | 1.03 | 1.02 | 0.98 | 1.00 | 0.98 | 1.04 | 1.00 | 1.02 | 1.01 | 1.01 | 0.96 | 0.97 | 1.13 | 0.75 |
| AG-c15_1 | Female | 6.84 | 0.86-1.14 ; 0.79-1.21 | 0.94 | 0.99 | 0.97 | 1.06 | 1.01 | 0.99 | 0.96 | 1.05 | 0.99 | 1.02 | 1.01 | 1.06 | 0.98 | 1.01 | 1.02 | 0.97 | 0.93 | 1.02 | 0.97 | 1.01 | 0.95 | 1.03 | 0.98 | 1.02 | 1.14 | 0.73 |
| AG-c15_2 | Female | 6.55 | 0.87-1.13 ; 0.80-1.20 | 0.99 | 0.96 | 0.98 | 1.06 | 0.98 | 1.04 | 1.04 | 1.00 | 0.99 | 1.02 | 0.99 | 1.05 | 0.96 | 1.03 | 1.07 | 0.93 | 0.98 | 0.96 | 1.06 | 1.02 | 0.96 | 0.98 | 0.90 | 1.01 | 1.13 | 0.71 |
| Oxr-c2_1 | Female | 6.08 | 0.88-1.12 ; 0.82-1.18 | 1.01 | 1.02 | 0.99 | 1.00 | 0.97 | 1.03 | 0.96 | 0.93 | 1.04 | 1.06 | 1.03 | 1.03 | 0.96 | 1.07 | 1.02 | 0.95 | 0.97 | 0.97 | 1.07 | 0.98 | 1.09 | 1.02 | 1.04 | 1.02 | 1.26 | 0.74 |
| Oxr-c2_2 | Female | 6.60 | 0.87-1.13 ; 0.80-1.20 | 0.93 | 1.01 | 0.99 | 1.02 | 1.00 | 0.99 | 0.92 | 0.96 | 1.04 | 0.99 | 1.05 | 1.06 | 0.91 | 1.08 | 1.01 | 0.96 | 0.99 | 0.99 | 1.07 | 0.98 | 1.04 | 0.97 | 1.02 | 0.96 | 1.20 | 0.77 |
| Oxr-c10_1 | Female | 6.73 | 0.87-1.13 ; 0.80-1.20 | 0.99 | 1.01 | 1.00 | 1.02 | 1.03 | 0.96 | 0.96 | 1.01 | 1.03 | 1.05 | 1.00 | 0.98 | 1.02 | 1.03 | 0.99 | 0.96 | 0.96 | 0.96 | 1.04 | 0.94 | 1.06 | 0.96 | 0.92 | 1.04 | 1.22 | 0.68 |
| Oxr-c10_2 | Female | 7.28 | 0.85-1.15 ; 0.78-1.22 | 0.97 | 1.00 | 1.00 | 0.99 | 1.04 | 0.98 | 0.95 | 0.96 | 1.05 | 0.98 | 0.99 | 0.97 | 1.05 | 1.07 | 1.00 | 0.99 | 0.97 | 0.94 | 1.04 | 0.95 | 1.12 | 0.96 | 0.93 | 1.02 | 1.19 | 0.78 |

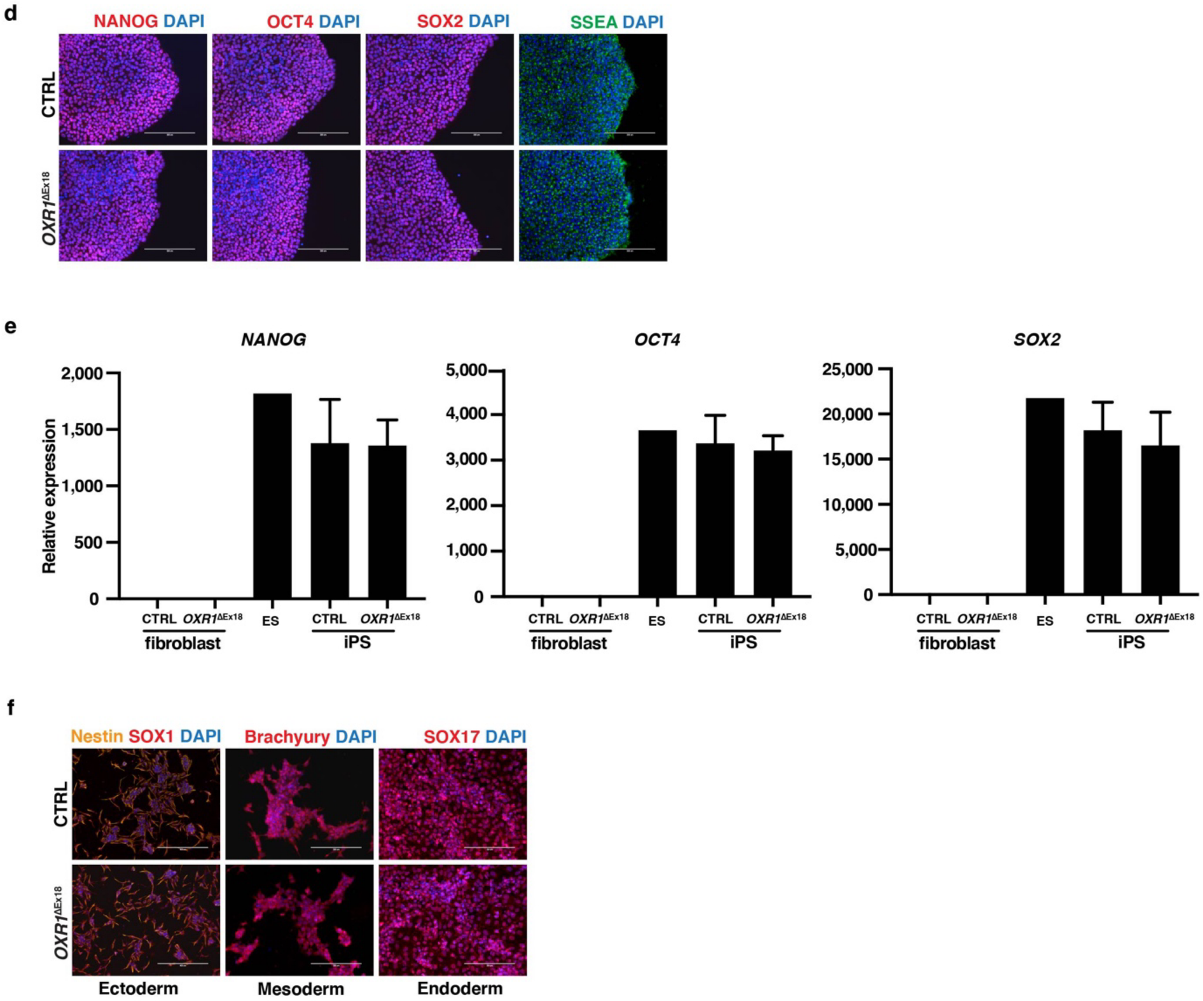

**g**

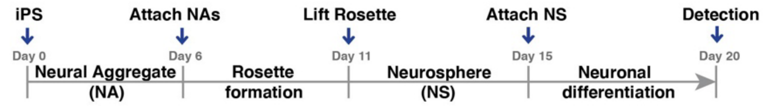
iPS
Attach NAs
Lift Rosette
Attach NS
Detection
Day 0
Day 6
Day 11
Day 15
Day 20
Neural Aggregate (NA)
Rosette formation
Neurosphere (NS)
Neuronal differentiation

**h**

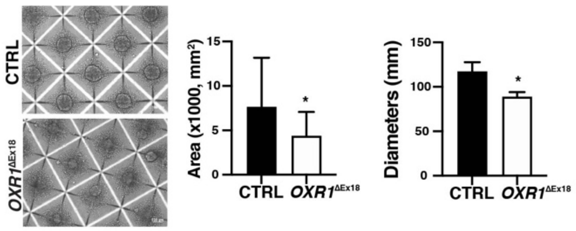
CTRL
OXR1ΔEx18
Area (x1000, mm²)
15
10
5
0
*
CTRL
OXR1ΔEx18
Diameters (mm)
150
100
50
0
*
CTRL
OXR1ΔEx18

**i**

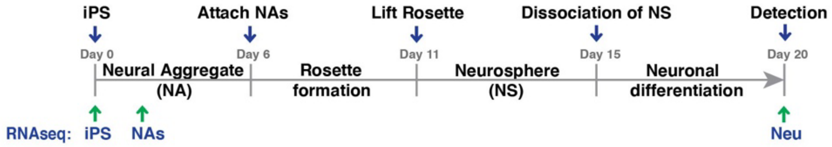
iPS
Attach NAs
Lift Rosette
Dissociation of NS
Detection
Day 0
Day 6
Day 11
Day 15
Day 20
Neural Aggregate (NA)
Rosette formation
Neurosphere (NS)
Neuronal differentiation
RNAseq: iPS
NAs
Neu

**j**

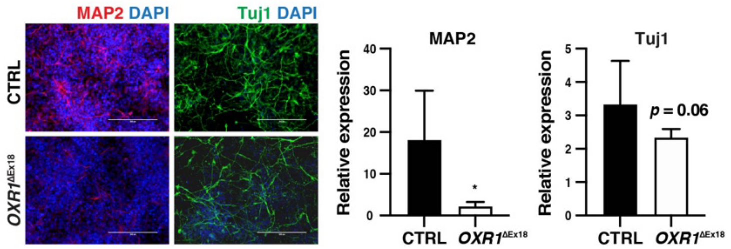
MAP2 DAPI
Tuj1 DAPI
CTRL
OXR1ΔEx18
MAP2
Relative expression
40
30
20
10
0
*
CTRL
OXR1ΔEx18
Tuj1
Relative expression
5
4
3
2
1
0
p = 0.06
CTRL
OXR1ΔEx18

**k**

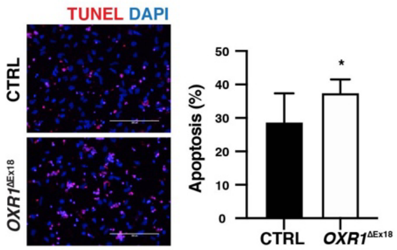
TUNEL DAPI
CTRL
OXR1ΔEx18
Apoptosis (%)
50
40
30
20
10
0
*
CTRL
OXR1ΔEx18

**l**

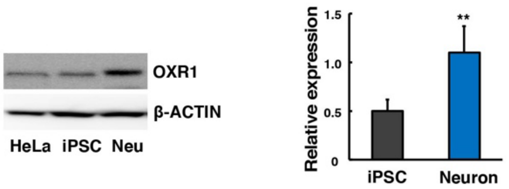
OXR1
β-ACTIN
HeLa
iPSC
Neu
Relative expression
1.5
1.0
0.5
0
**
iPSC
Neuron

m

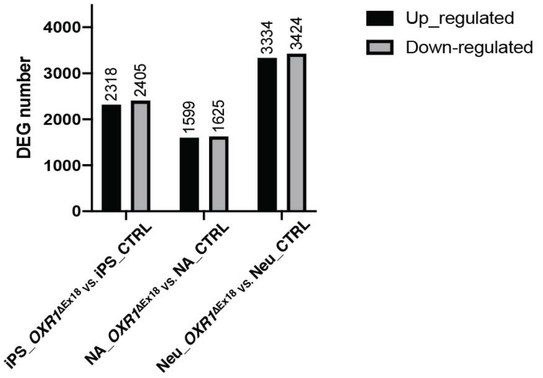
DEG number
4000
3000
2000
1000
0
2318
2405
1599
1625
3334
3424
Up_regulated
Down-regulated
iPS_OXR1ΔEx18 vs. iPS_CTRL
NA_OXR1ΔEx18 vs. NA_CTRL
Neu_OXR1ΔEx18 vs. Neu_CTRL

n

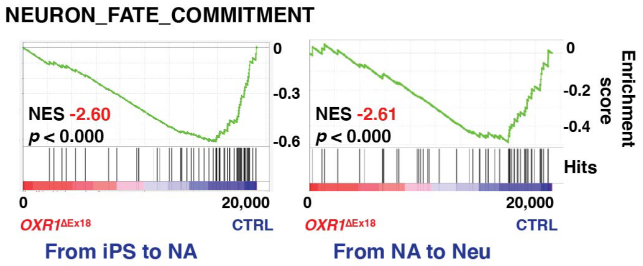
NEURON_FATE_COMMITMENT
NES -2.60
p < 0.000
0
-0.3
-0.6
0
20,000
OXR1ΔEx18
CTRL
From iPS to NA
NES -2.61
p < 0.000
0
-0.2
-0.4
Enrichment score
Hits
0
20,000
OXR1ΔEx18
CTRL
From NA to Neu

o

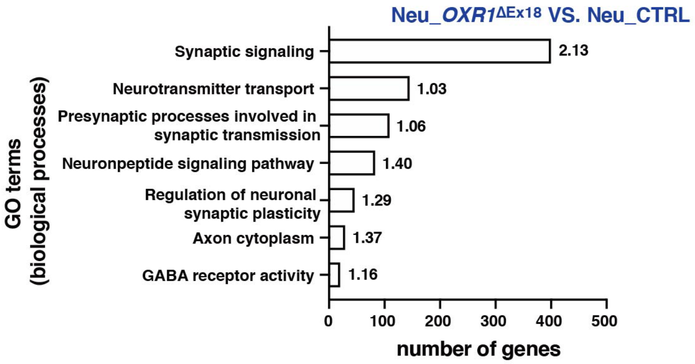
Neu_OXR1ΔEx18 VS. Neu_CTRL
GO terms (biological processes)
Synaptic signaling
2.13
Neurotransmitter transport
1.03
Presynaptic processes involved in synaptic transmission
1.06
Neuronpeptide signaling pathway
1.40
Regulation of neuronal synaptic plasticity
1.29
Axon cytoplasm
1.37
GABA receptor activity
1.16
0
100
200
300
400
500
number of genes

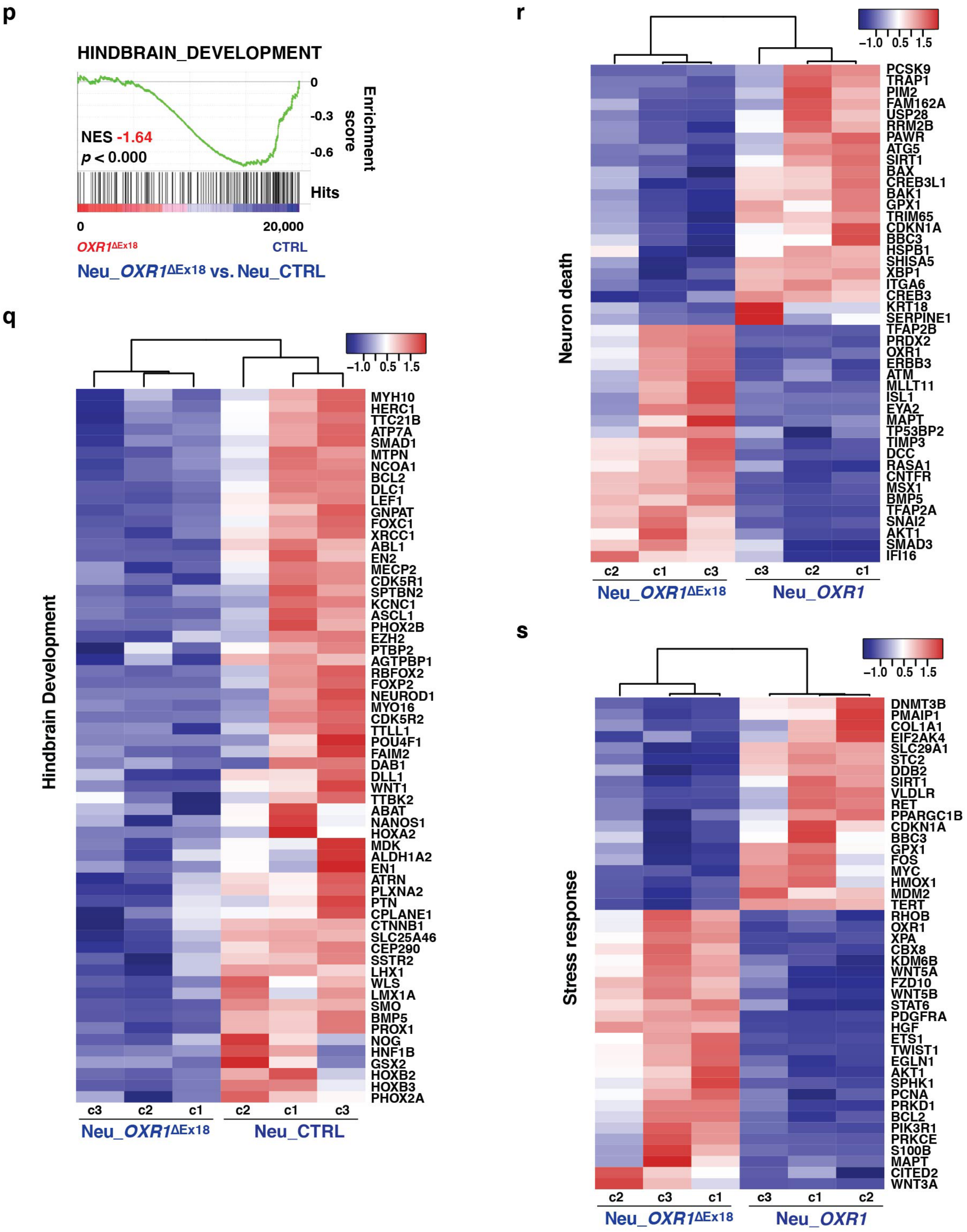
p
HINDBRAIN_DEVELOPMENT
NES -1.64
p < 0.000
0
-0.3
-0.6
Enrichment score
Hits
0
20,000
OXR1ΔEx18
CTRL
Neu_OXR1ΔEx18 vs. Neu_CTRL
q
−1.0 0.5 1.5
Hindbrain Development
MYH10
HERC1
TTC21B
ATP7A
SMAD1
MTPN
NCOA1
BCL2
DLC1
LEF1
GNPAT
FOXC1
XRCC1
ABL1
EN2
MECP2
CDK5R1
SPTBN2
KCNC1
ASCL1
PHOX2B
EZH2
PTBP2
AGTPBP1
RBFOX2
FOXP2
NEUROD1
MYO16
CDK5R2
TTLL1
POU4F1
FAIM2
DAB1
DLL1
WNT1
TTBK2
ABAT
NANOS1
HOXA2
MDK
ALDH1A2
EN1
ATRN
PLXNA2
PTN
CPLANE1
CTNNB1
SLC25A46
CEP290
SSTR2
LHX1
WLS
LMX1A
SMO
BMP5
PROX1
NOG
HNF1B
GSX2
HOXB2
HOXB3
PHOX2A
c3 c2 c1 c2 c1 c3
Neu_OXR1ΔEx18
Neu_CTRL
r
−1.0 0.5 1.5
Neuron death
PCSK9
TRAP1
PIM2
FAM162A
USP28
RRM2B
PAWR
ATG5
SIRT1
BAX
CREB3L1
BAK1
GPX1
TRIM65
CDKN1A
BBC3
HSPB1
SHISA5
XBP1
ITGA6
CREB3
KRT18
SERPINE1
TFAP2B
PRDX2
OXR1
ERBB3
ATM
MLLT11
ISL1
EYA2
MAPT
TP53BP2
TIMP3
DCC
RASA1
CNTFR
MSX1
BMP5
TFAP2A
SNAI2
AKT1
SMAD3
IFI16
c2 c1 c3 c3 c2 c1
Neu_OXR1ΔEx18
Neu_OXR1
s
−1.0 0.5 1.5
Stress response
DNMT3B
PMAIP1
COL1A1
EIF2AK4
SLC29A1
STC2
DDB2
SIRT1
VLDLR
RET
PPARGC1B
CDKN1A
BBC3
GPX1
FOS
MYC
HMOX1
MDM2
TERT
RHOB
OXR1
XPA
CBX8
KDM6B
WNT5A
FZD10
WNT5B
STAT6
PDGFRA
HGF
ETS1
TWIST1
EGLN1
AKT1
SPHK1
PCNA
PRKD1
BCL2
PIK3R1
PRKCE
S100B
MAPT
CITED2
WNT3A
c2 c3 c1 c3 c1 c2
Neu_OXR1ΔEx18
Neu_OXR1

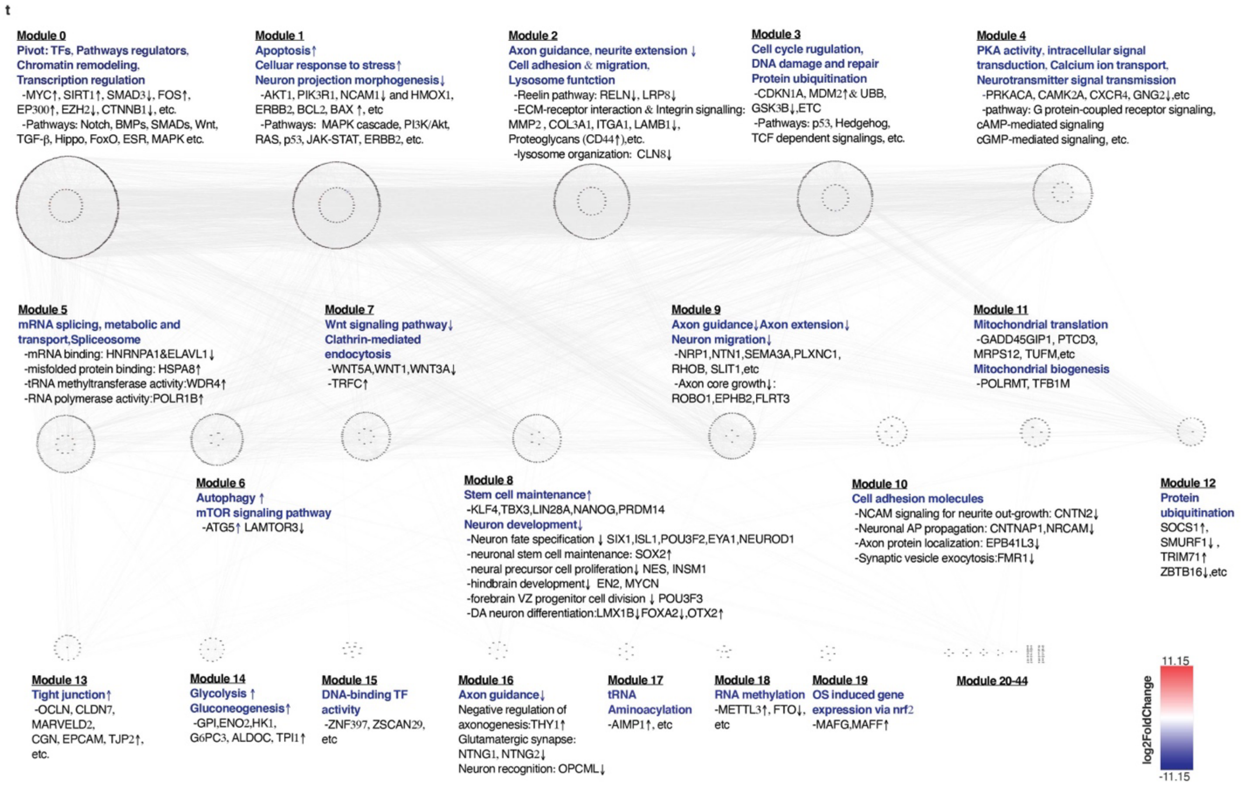


**Supplementary Figure 3. Generation and characterization of 2D neuronal differentiation models derived from induced pluripotent stem cells. a,** Genotyping of generated iPS from healthy control clones (CTRL) and OXR1-deficient iPS clones (*OXR1*$^{\Delta Ex18}$). After total RNA and cDNA were prepared, PCR was performed using primer spanning exon 18 region. The shorter (185bp) PCR products were detected in all three *OXR1*$^{\Delta Ex18}$ iPSC clones. **b,** Immunoblot of OXR1 in CTRL and *OXR1*$^{\Delta Ex18}$ iPS showing complete lack of OXR1 protein in patient sample. **c,** Karyotype analysis of fibroblast cells and iPSC clones. Only qualified iPS clones were selected in further experiments. **d,** Immunofluorescence of pluripotent transcription factors OCT4, SOX2, NANOG, and cell surface marker SSEA4 in iPS. **e,** Gene expression of endogenously expressed pluripotency transcription factors in both iPS and fibroblasts from CTRL and *OXR1*$^{\Delta Ex18}$ by qPCR analysis. The gene expression levels were comparable with human embryonic stem (ES) cells. **e,** iPS were differentiated into the three germ layers *in vitro* by using STEMdiff™ Trilineage Differentiation Kit. Immunocytochemistry was performed to identify the differentiation of ectodermal (SOX1 and Nestin), mesodermal (Brachyury), or endodermal (SOX17) lineages. **g,** Schematic representation of experimental procedure for neurite outgrowth. **h**, Phase contrast images of NAs generated in AggreWell at 48 h after aggregation (left) showing irregularities in the morphology of *OXR1*$^{\Delta Ex18}$. Quantification of diameters (middle) and areas (right) of NAs at 96 h after aggregation determined by ImageJ. Four hundred cells of each group from two independent cultures were counted. **i**, Schematic representation of experimental procedure for neuronal differentiation and RNA sequence (RNAseq). Neu, neuron. **j**, Immunofluorescence images(left) and quantifications(right) of the fluorescence intensity of MAP2 and Tuj1 from CTRL and $OXR1^{\Delta Ex18}$. The scale bars, 200 μm. **k**, Representative images of TUNEL staining of apoptotic cells during neural differentiation (left) and quantification of TUNEL staining in control and $OXR1^{\Delta Ex18}$ neuronal cells (right). About 1,300 cells of each group from 8 different microscopic fields were counted. The scale bars, 200 μm. **l**, Western blot images(right) and quantifications(left) of OXR1 expression, Neuron vs. undifferentiated iPS, n=3. **h**, **j- l,** n=3, Data are means+SD, * $p < 0.05$, ** $p < 0.01$, *t*-test. **m**, Bar chart showing the number of Down-regulated and Up-regulated differentially expressed genes (DEGs) between patient ($OXR1^{\Delta Ex18}$) and control (CTRL) at each differentiation stage. Thresholds at logFC ≥ 0.5 (Down-DEGs) and ≤ -0.5 (Up-DEGs), significant at adjusted-*p* value < 0.05. **n**, Gene set enrichment analysis (GSEA) showing significant reduction in neuron fate commitment following neuronal differentiation in $OXR1^{\Delta Ex18}$ groups when compared to CTRL. NES, normalized enrichment score; *p*, nominal *p* value. **o**, Bar charts of the top ranked (upregulated) GO Terms of biological processes relevant to neural development, between $OXR1^{\Delta Ex18}$ and CTRL at neuron stage, identified by GSEA. **p**, Impaired hindbrain development in $OXR1^{\Delta Ex18}$ neurons. GSEA showing downregulated of hindbrain development in Neu_$OXR1^{\Delta Ex18}$, as compared with Neu_CTRL. **q**, Heatmaps of leading-edge genes of hindbrain development by GSEA in the comparison of Neu_$OXR1^{\Delta Ex18}$ vs. Neu_CTRL. **r-s**, Heatmaps of top 40 leading-edge genes related to neuron death(**r**) or stress response(**s**) in the comparison of Neu_ $OXR1^{\Delta Ex18}$ vs. Neu_CTRL. **t**, Topologic biological network analysis of the protein and protein interactions of 1539 DEGs between Neu_$OXR1^{\Delta Ex18}$ and Neu_CTRL. Each module consists of a set of genes that are both highly connected in protein interaction networks and highly correlated in biological databases and group in a circle. For instance, Module 0 functioned as a pivot for the whole network characterized by central transcription factors, binding proteins, and core enzymatic components (HDAC1, SIRT1, EP300, EZH2, etc) crucial for regulating transcription, chromatin organization and several substantial pathways for cellular activities (Notch, BMPs, SMADs and Wnt, etc). Coordinated with Module 1 and 3, Module 0 regulated cell stress responses. Module 8 conferred a lead role in neurogenesis. Module 2, 9, 10 and 16 showed that OXR1-depletion impaired axon guidance at multiple levels, from the axon genesis, axon core growth, to neurite extension, migration, recognition, then to synapse formation.The boundary color of a node(gene) represents its module. Node size represents "degree" describing the relative importance of a single node within the sub-network. Hub genes, highly connected nodes, lie in the inner circle of each module identified by Cytohubba. Node color reflects whether its mRNA is up or down regulated in Neu_$OXR1^{\Delta Ex18}$ in comparison with Neu_CTRL. Color descending from red to blue, indicates log2 (Fold Change) value from large to small. Visualization by Cytoscape 3.8.1. iPS, induced pluripotent stem cell; NA, neural aggregates; Neu, neurons.

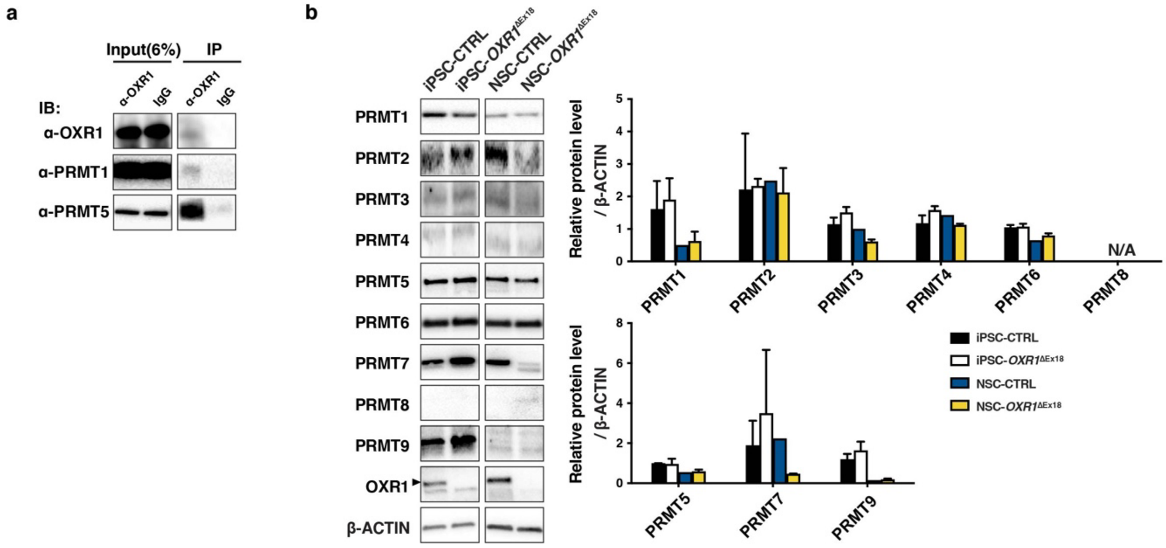


**Supplementary Figure 4. OXR1 interacts with PRMT1 and PRMT5. a,** Immunoprecipitation of OXR1 in U2OS cells demonstrating that OXR1 bound to PRMT1 and PRMT5 respectively. 6% of extracted protein was loaded as input control. **b**, Western blot analysis of PRMTs in iPS and NSC. Quantifications of type I (upper)and type II/III (down)of PRMTs. β-ACTIN was used as loading control. Results are shown as means+SD, *t*-test. iPS, induced pluripotent stem cells; NSC, neural stem cells.

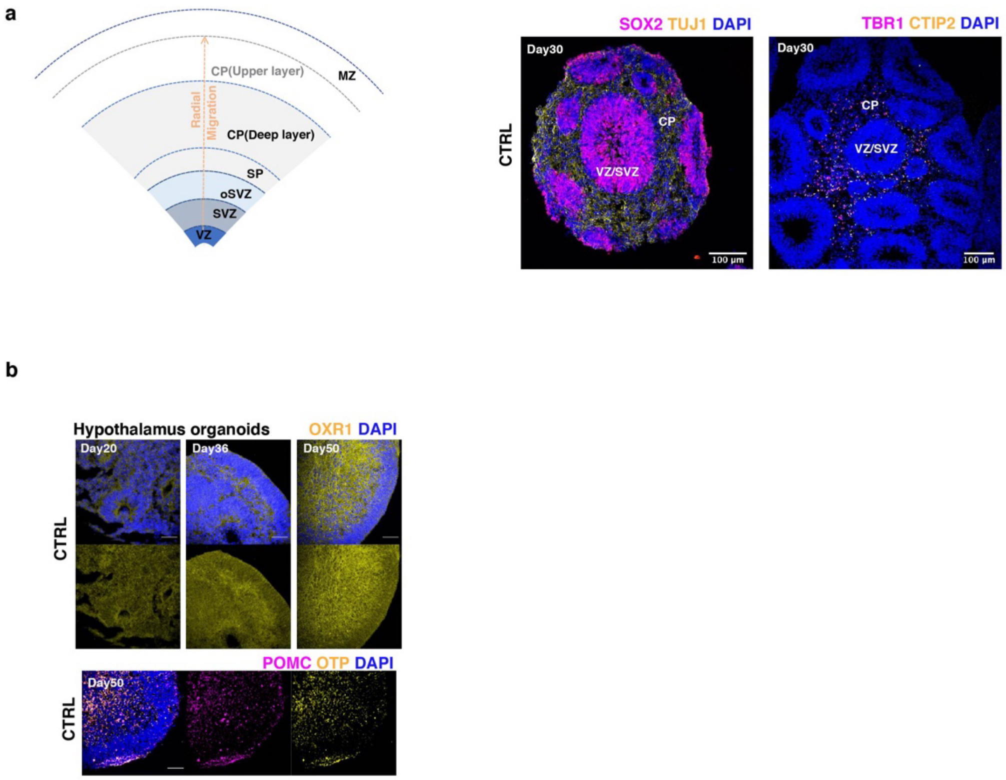


**Supplementary Figure 5. Generation and characterization of cerebral organoids and hypothalamus organoids derived from iPS. a,** A schematic of cortical layer structure of cerebral organoids or cortical organoids at ≥Day 75(left). The dotted arrow in orange indicating the radial migration of neural progenitor cells from VZ/SVZ. The SP like region is usually interspersed between SVZ and CP. Representative confocal images of cortical region at Day30 (right). The progenitor zone consists of VZ and SVZ, stained with SOX2. TBR1 and CITP2 marked the deep layer neurons of CP, and the nuclei were visualized using DAPI. VZ, ventricular zone; SVZ, subventricular zone; oSVZ, outer SVZ; SP, subplate; CP, cortical plate ; MZ, marginal zone. **b,** Representative tiling confocal images showing the development of hypothalamus organoids of CTRL immunostained with anti-OXR1 antibodies at Days 20, 35, and 50(upper). OXR1 level increased along with neuronal maturation. Representative tiling confocal images showing control hypothalamus organoids contained POMC$^+$ peptidergic neurons and OTP$^+$ hypothalamic neurons at Day50 (bottom).

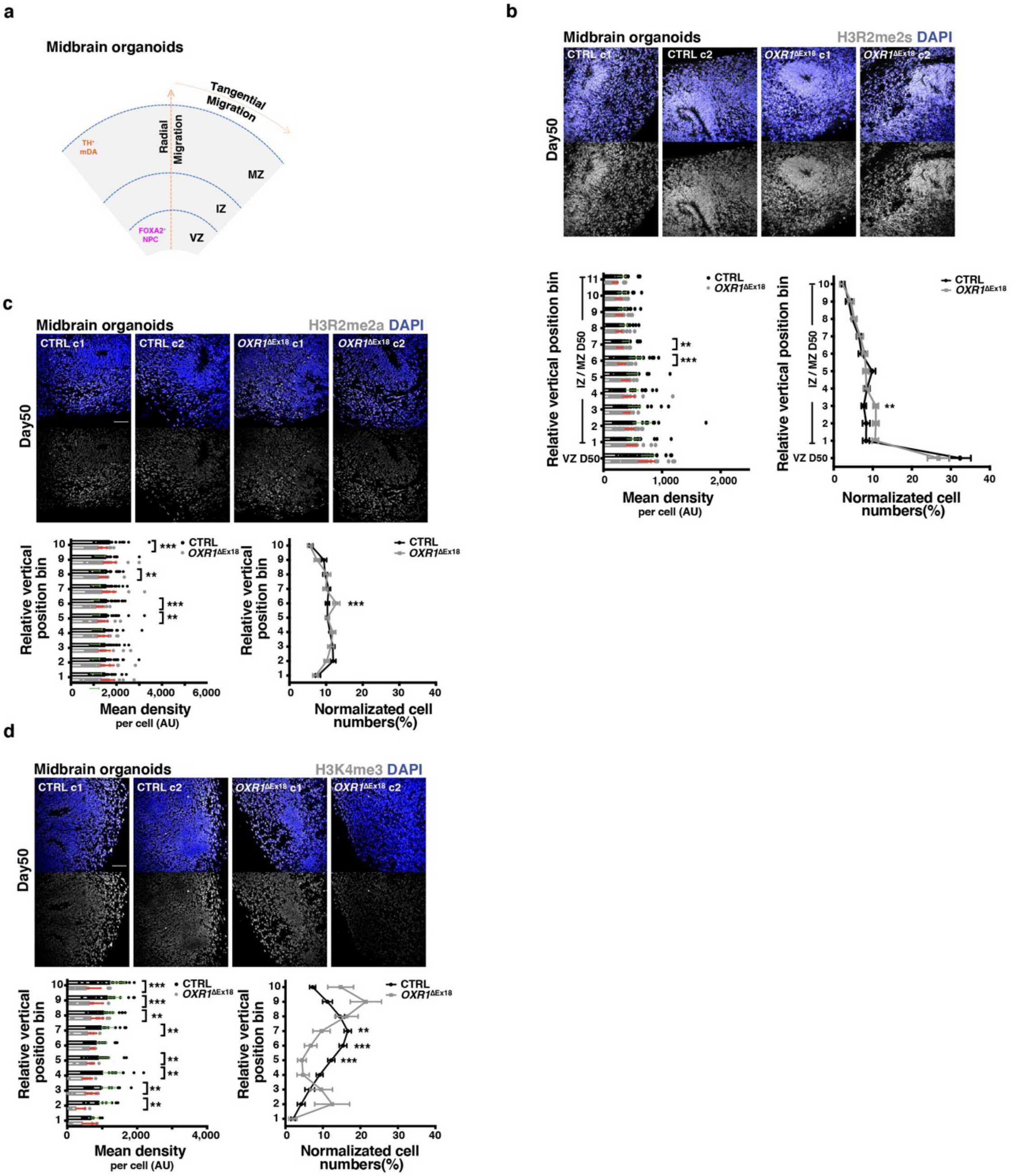


**Supplementary Figure 6. OXR1 impacts midbrain development by shaping histone arginine and lysine methylations. a**, A schematic of midbrain organoid, the dotted arrow in orange indicating the radial migration of neural progenitor cells from VZ, through IZ, to reach MZ, and tangential migration of $TH^{+}$ neurons at the outer surface of MZ. **b-d**, Representative images, and quantifications of the distribution of $H3R2me2s^{+}$ (**b**), $H3R2me2a^{+}$(**c**), and $H3K4me3^{+}$ (**d**) cells in midbrain organoids at Day50. The dense radial developing structures from apical surface are labelled as VZ. The outside VZ structure (IZ/MZ) from the outer boundary of VZ to basal surface is evenly divided into 10 bins, from 1 to 10. Quantifications include the average expression level per cell(lower left) and the normalized abundance within each bin and in VZ(lower right) . The normalized abundance was calculated as (the number of positive cells in a bin/ the total number of positive cells). Values represent mean ± SEM, ***p* < 0.01; ****p* < 0.001; Student's *t* test. In H3R2me2s$^{+}$ (**b**), CTRL, n = 23; *OXR1*$^{\Delta Ex18}$, n =33, from two independent batches; in H3R2me2a$^{+}$(**c**), CTRL, n = 31; *OXR1*$^{\Delta Ex18}$, n = 21, from two independent batches; in H3K4me3$^{+}$ (**d**), CTRL, n = 31; *OXR1*$^{\Delta Ex18}$, n = 21, from two independent batches. VZ, ventricular zone; IZ, intermediate zone; MZ, mantle zone, NPC, neural progenitor cells.

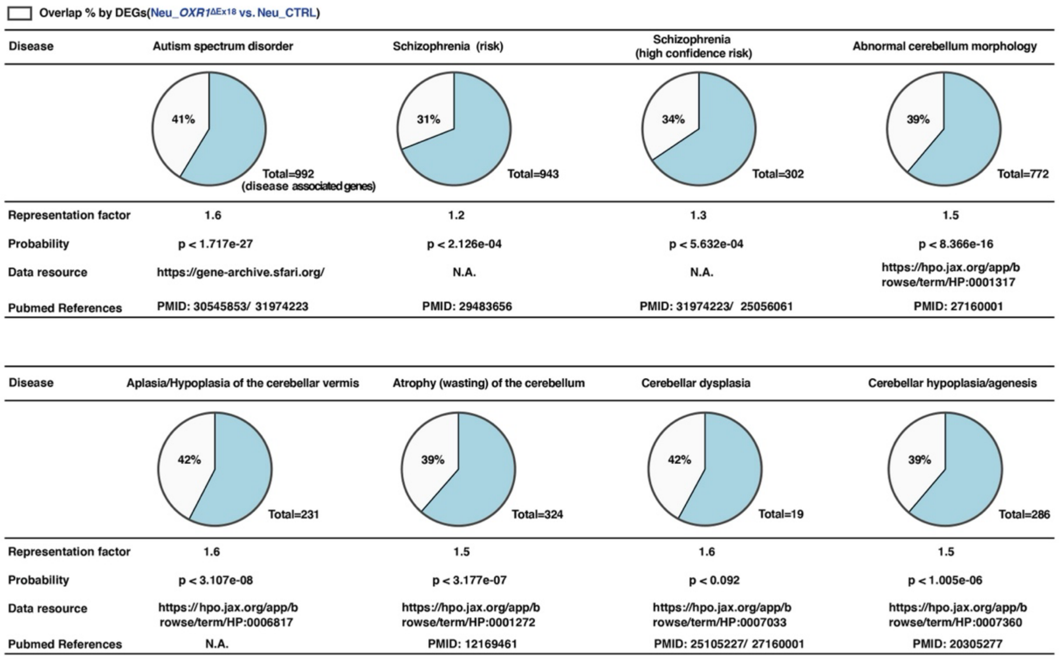


**Supplementary Figure 7. Intersection-comparisons of OXR1 deficiency disease and other neurological developmental diseases.** The representation factors and relevant probability are calculated by http://nemates.org/MA/progs/overlap_stats.html . A representation factor > 1 indicates more overlap than expected of two independent gene sets, the set of differentially expressed genes (DEGs) of Neu_ *OXR1*$^{\Delta Ex18}$ vs. Neu_CTRL and the disease associated gene set of the neurological developmental disease as indicated.
